# Mechanisms of USP7/MAGEL2 Complex Assembly and Its Mutational Disruption in Neurodevelopmental Diseases

**DOI:** 10.64898/2026.04.24.720667

**Authors:** Emilie J. Korchak, Gabriella A. Soriano, Irina Semenova, Bing Hao, Denis Štepihar, Tara Bayat, Klementina Fon Tacer, Irina Bezsonova

## Abstract

The WASH complex regulates endosomal trafficking and is linked to several neurodevelopmental diseases, including Prader-Willi syndrome, Schaaf-Yang syndrome, and Hao-Fountain syndrome. Its function is tightly controlled by ubiquitination, maintained by the multi-subunit MUST complex containing both a ubiquitin ligase (MAGEL2/TRIM27) and a deubiquitinase (USP7). However, the mechanism underlying the MUST complex assembly remains poorly understood. In this study, we investigate the assembly of USP7 and MAGEL2 components of the MUST complex using NMR spectroscopy, isothermal titration calorimetry, X-ray crystallography, and cellular assays. We show that the USP7/MAGEL2 interaction is bipartite and multivalent. Two distinct domains of USP7, TRAF and UBL1-2, recognize two unstructured but evolutionarily conserved regions of MAGEL2, one of which contains multiple TRAF-binding sites. Furthermore, we determine the high-resolution crystal structure of the TRAF/MAGEL2 complex and identify Hao-Fountain syndrome-linked mutations in USP7 that disrupt USP7/MAGEL2 complex formation *in vitro* and in cells. These findings provide mechanistic insight into the pathogenic basis of Hao-Fountain syndrome and related Schaaf-Yang and Prader-Willi syndromes.

## INTRODUCTION

Ubiquitination is a reversible post-translational modification that regulates protein degradation, signal transduction, gene expression, and DNA repair. The ubiquitin-mediated signaling is essential for the maintenance of protein homeostasis^1,2^. Consequently, defects in enzymes responsible for adding and removing ubiquitin from the target protein result in diseases, including cancers, neurogenetic and neurodegenerative disorders, and immune system dysfunctions^3-6^. The WASH complex (Wiskott Aldrich Syndrome protein and SCAR homologue complex) regulates endosomal trafficking, and its function is tightly controlled by ubiquitination, in which K63-linked ubiquitin chains are added to and removed from the complex through the coordinated actions of the E3 ubiquitin ligase complex MAGEL2/TRIM27 and its counterpart, the deubiquitinating enzyme USP7. TRIM27 ubiquitinates WASH, thereby activating it to promote endosomal actin assembly and recycling^7-10^, while USP7 acts as a molecular rheostat that deubiquitinates both the WASH complex and TRIM27, balancing their activity and stability^11^. MAGEL2 bridges TRIM27 and USP7, forming a heterotrimeric multifunctional MUST complex (<u>M</u>AGEL2/<u>US</u>P7/<u>T</u>RIM27) that regulates WASH-mediated endosomal protein recycling^9,12,13^. This precise interplay between ubiquitination and deubiquitination ensures proper protein distribution and function during cellular differentiation and tissue development.

Defects in MAGEL2 and USP7 have been linked to several rare neurodevelopmental disorders with overlapping phenotypes. Deletions of the MAGEL2-coding gene result in Prader-Willi syndrome, while its truncations cause Schaaf-Yang syndrome, both of which present with neonatal hypotonia which turns into hyperphagia later in life, hormonal disbalance, and developmental delays with intellectual disability and autism spectrum disorder^14-19^. Deletions and mutations of the *USP7* gene lead to another neurodevelopmental disorder known as Hao-Fountain syndrome, which shares clinical features with Prader-Willi and Schaaf-Yang syndromes, including hypogonadism, hypotonia, and intellectual disability^11,20^. These related disorders underscore the significance of the WASH protein complex in regulating protein recycling during development and highlight the need for a better understanding of the mechanisms governing its activation through MUST complex formation.

MAGEL2 is a member of a family of approximately 40 proteins that fall into two main groups: cancer-testis antigens, and proteins enriched in the brain^21^. MAGEL2 is expressed predominantly in the hypothalamus^22,23^ and consists of a large, intrinsically disordered, N-terminal region and a C-terminal MAGE Homology Domain (MHD), conserved in all MAGEs (**Figure 1A**). The disordered region is necessary for engaging USP7^11^. USP7 contains a catalytic domain, a TRAF domain and five ubiquitin-like domains (UBL1-5)^24,25^. The TRAF and UBL1-2 domains serve as the primary substrate-binding modules that ensure specificity toward diverse substrates^26,27^, and the Hao-Fountain syndrome patient mutations analyzed in this study cluster within these domains.

**Figure 1.**
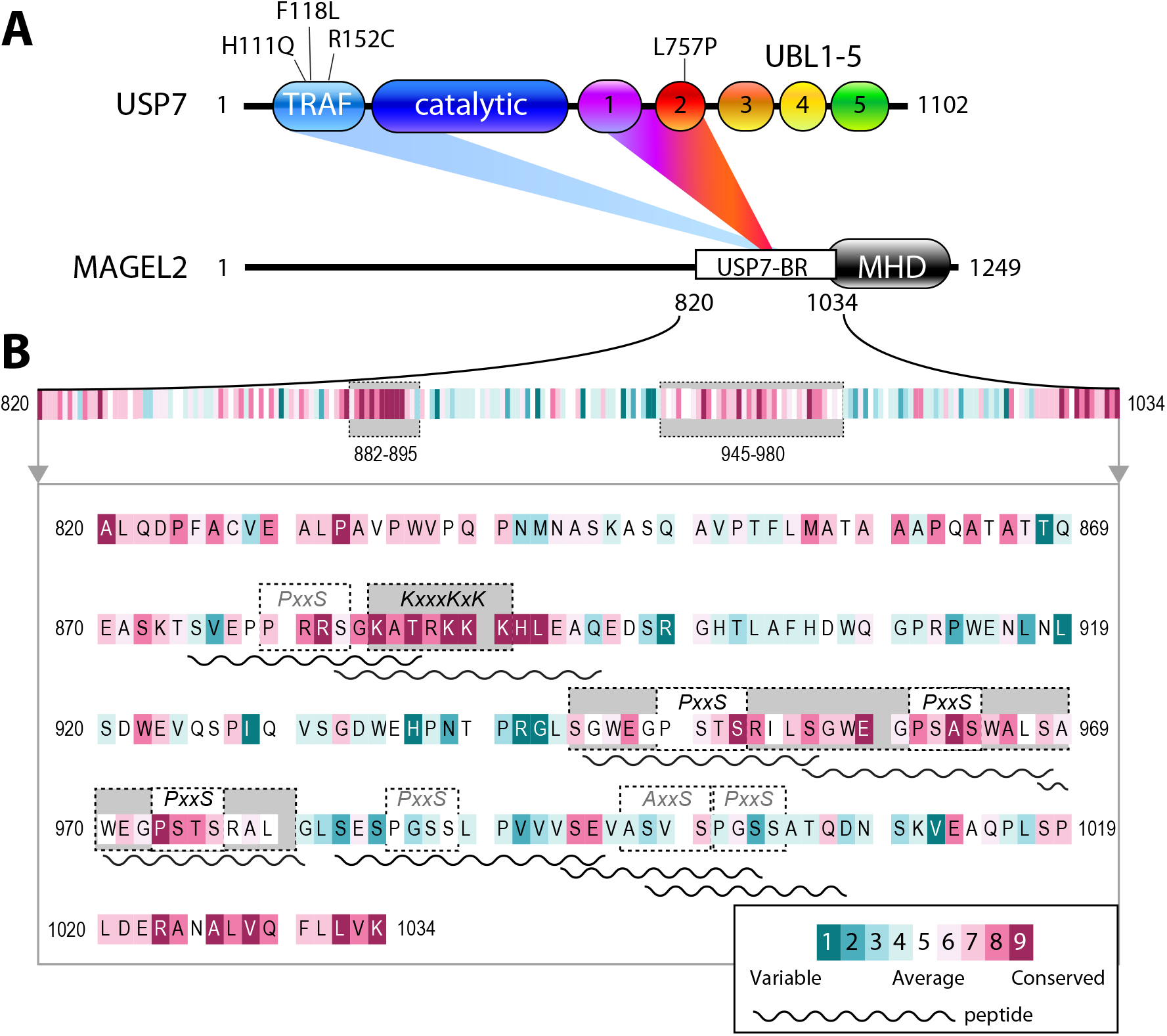
MAGEL2 contains multiple USP7-binding motifs. **A**. Schematic of USP7 and MAGEL2, highlighting key structural features. USP7 domains include the TRAF domain (light blue), catalytic domain (dark blue), and ubiquitin-like domains 1-5 (UBL1 purple, UBL2 red, UBL3 orange, UBL4 yellow, UBL5 green). The Hao-Fountain syndrome-linked missense mutations in the TRAF and UBL1-2 domains are indicated. MAGEL2 shows the MAGE homology domain (MHD, black) and the disordered USP7 binding region (USP7-BR, white). **B**. Zoomed-in view of the USP7 binding region (residues 820-1034) in MAGEL2, showing potential USP7-binding motifs (boxed). Synthetic peptides corresponding to MAGEL2 regions are represented as wavy lines. Amino acid residues are color-coded by conservation score, ranging from teal (highly variable) to magenta (highly conserved).

Many USP7 targets are recognized by the TRAF domain^28-33^, while others interact exclusively with UBL1-2^25,34-37^. A few substrates simultaneously engage both the TRAF and UBL1-2 domains, with TRAF serving as the primary binding site, as demonstrated for FBXO38^38^, PAF15^39^, and DNA Polymerase iota^40^. These observations suggest that USP7 employs a modular and potentially multivalent mode of substrate recognition, raising the possibility that similar mechanisms may underlie its interaction with MAGEL2.

Despite the established roles of USP7 and MAGEL2 in the MUST complex, the molecular basis of their interaction and the effects of patient-derived mutations on complex assembly remain largely unexplored. In this study, we pinpoint functional USP7-binding motifs in MAGEL2 and characterize USP7/MAGEL2 complexes using complementary biochemical, structural, and cellular approaches. NMR spectroscopy and X-ray crystallography reveal how the TRAF and UBL1-2 domains engage distinct regions of MAGEL2 with comparable affinities, highlighting a bipartite mode of substrate recognition. We further identify pathogenic USP7 variants that disrupt complex formation both *in vitro* and in cells. Together, these findings provide mechanistic and structural insights into USP7 substrate recognition and explain how disease-linked mutations compromise this process, contributing to the molecular pathology of Hao-Fountain syndrome and related MUST complex disorders.

## RESULTS

### MAGEL2 harbors multiple functional USP7-binding motifs

Human MAGEL2 contains a large intrinsically disordered region that mediates its interaction with USP7 (**Figure 1A**). Previous studies have shown that residues 820-1034 within this region are required for efficient recruitment of USP7 to the MUST complex *in vitro* and in cells^11^. To identify specific sites driving MAGEL2/USP7 complex formation, we analyzed residues 820-1034 of MAGEL2 for canonical USP7-binding motifs, including E/A/PxxS^33^ and KxxxKxK/R^34^ sequences recognized by the TRAF and UBL1-2 domains of USP7, respectively. This analysis revealed seven putative TRAF-binding motifs and a single predicted UBL1-2-binding motif (**Figure 1B**, boxed).

To validate these interactions and determine which motifs preferentially bind USP7, we chemically synthesized peptides corresponding to each site (**Figure 1B**, wavy lines) and studied their binding using NMR chemical shift perturbation (CSP) assays (**Supplementary Figure S1**). Specifically, isotopically ^15^N-labeled TRAF or UBL1-2 domains were titrated with unlabeled peptides containing corresponding USP7-binding motifs, and the resulting CSPs in their NMR spectra were used to compare their affinities.

Of the seven potential TRAF-binding peptides tested, six MAGEL2 peptides 945-956, 956-968, 968-980, 982-995, 994-1003, and 998-1008 induced significant changes in their NMR spectra, confirming binding. In contrast, peptide 875-886 exhibited minimal CSPs, indicating a lack of interaction with TRAF (**Supplementary Figure S1A-G**). Additionally, the MAGEL2 region 882-895, containing a single putative highly conserved UBL1-2-binding motif KATRKKK, bound efficiently to the UBL1-2 (**Supplementary Figure S1H**). Among the six TRAF-binding peptides, three (945-956, 956-968, and 968-980) are evolutionarily conserved within this disordered region of MAGEL2, underscoring their likely functional significance. Accordingly, these peptides were selected for further study (**Figure 1B**).

### Validation of USP7/MAGEL2 interactions with isothermal titration calorimetry

NMR-detected binding was further confirmed with an orthogonal isothermal titration calorimetry (ITC) assay. All four binders chosen to further study (882-895, 945-956, 956-968, and 968-980) exhibit comparable dissociation constants in the low micromolar range as measured by ITC (**Figure 2**). TRAF-binding peptides 945-956, 956-968 and 968-980 bound to USP7 with 2.33±0.60 μM, 4.93±2.16 μM, and 1.29±0.55 μM affinities, respectively, in agreement with low micromolar affinities typically observed for TRAF-binding substrates^29,31-33,41,42^. The UBL1-2-binding peptide exhibited a K_D_ of 7.97± 2.81 μM.

**Figure 2.**
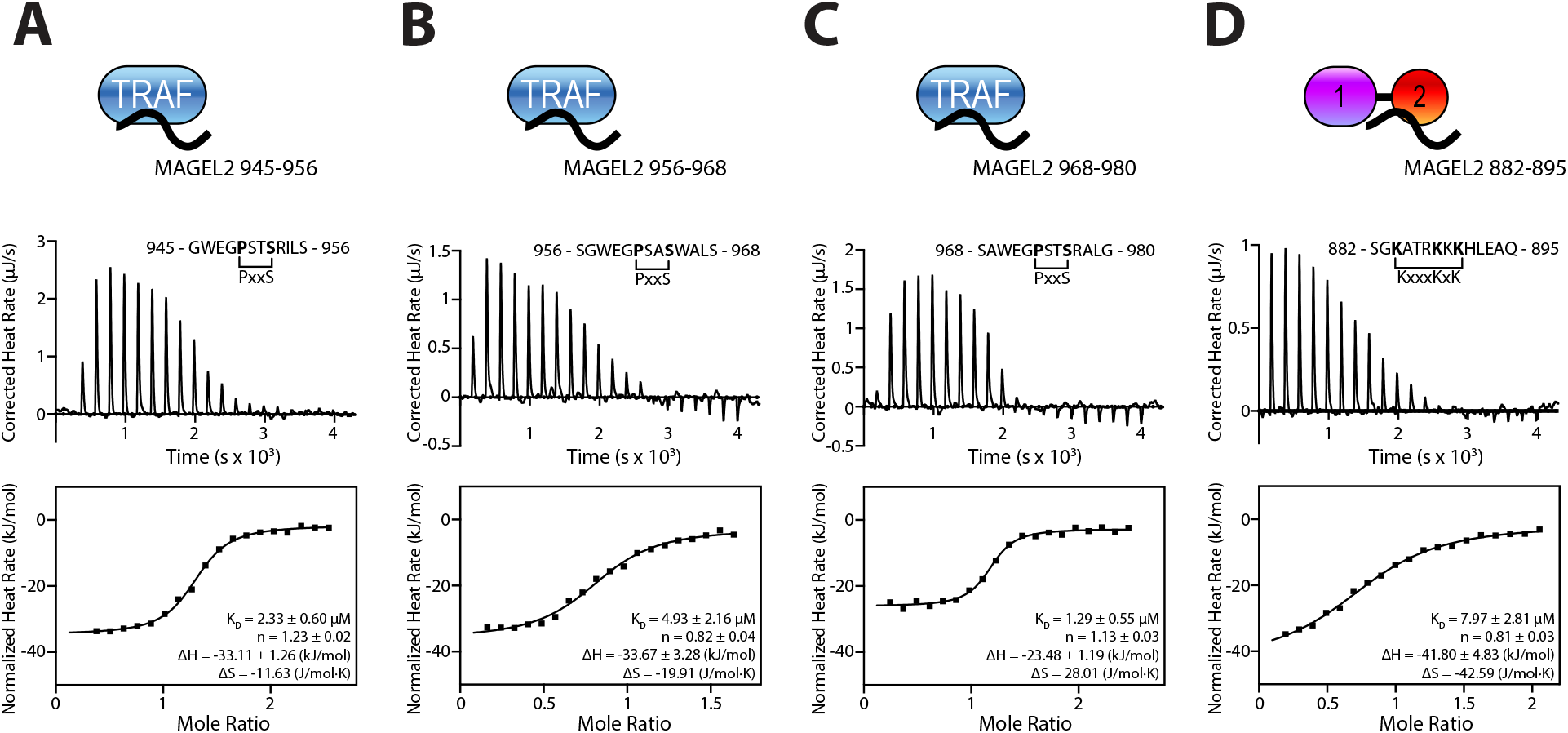
MAGEL2 binds to both TRAF and UBL1-2 domains of USP7. ITC binding isotherms of the USP7 TRAF domain titrated with MAGEL2 peptides (**A**) 945-956, (**B**) 956-968, and (**C**) 968-980, and of the USP7 UBL1-2 titrated with MAGEL2 882-895 (**D**).

Together, these findings demonstrate that USP7 recognizes MAGEL2 through a true bipartite interaction, engaging two distinct, evolutionarily conserved interfaces with similar affinities. This recognition mode is distinct from that of other USP7 substrates, such as Mdm2 and ICP0, which interact exclusively with either the TRAF or UBL1-2 domains, respectively^29,33,34,43^. Although FBXO38, PAF15 and DNA polymerase iota engage both domains, they do so with markedly asymmetric affinities, strongly favoring the TRAF domain^38,40^.

### NMR characterization of USP7/MAGEL2 binding interfaces

To determine the binding interfaces of TRAF/MAGEL2 complexes, we took advantage of the NMR titration data and measured the NMR chemical shift differences (Δω) between the free and MAGEL2-bound TRAF domain observed for each backbone amide peak of TRAF in its ^15^N-^1^H HSQC spectra (**Figure 3**). The mapping of the Δω magnitude on the surface of the TRAF domain revealed a similar binding interface for all three MAGEL2 peptides that includes residues F118, C121, and D164, which are located on the beta strands 4 and 7 of the TRAF, the classical substrate-recognition site of USP7^29-33,42,44^. Some of the central interface residues, such as W165, G166, and S168, could not be included in the analysis because their corresponding peaks are broadened beyond detection in the NMR spectrum, indicative of their conformational dynamics that occur on the µs-ms timescale. Nonetheless, the overall mapping clearly indicates that the TRAF domain of USP7 interacts with all three peptides *via* its canonical substrate-binding interface.

**Figure 3.**
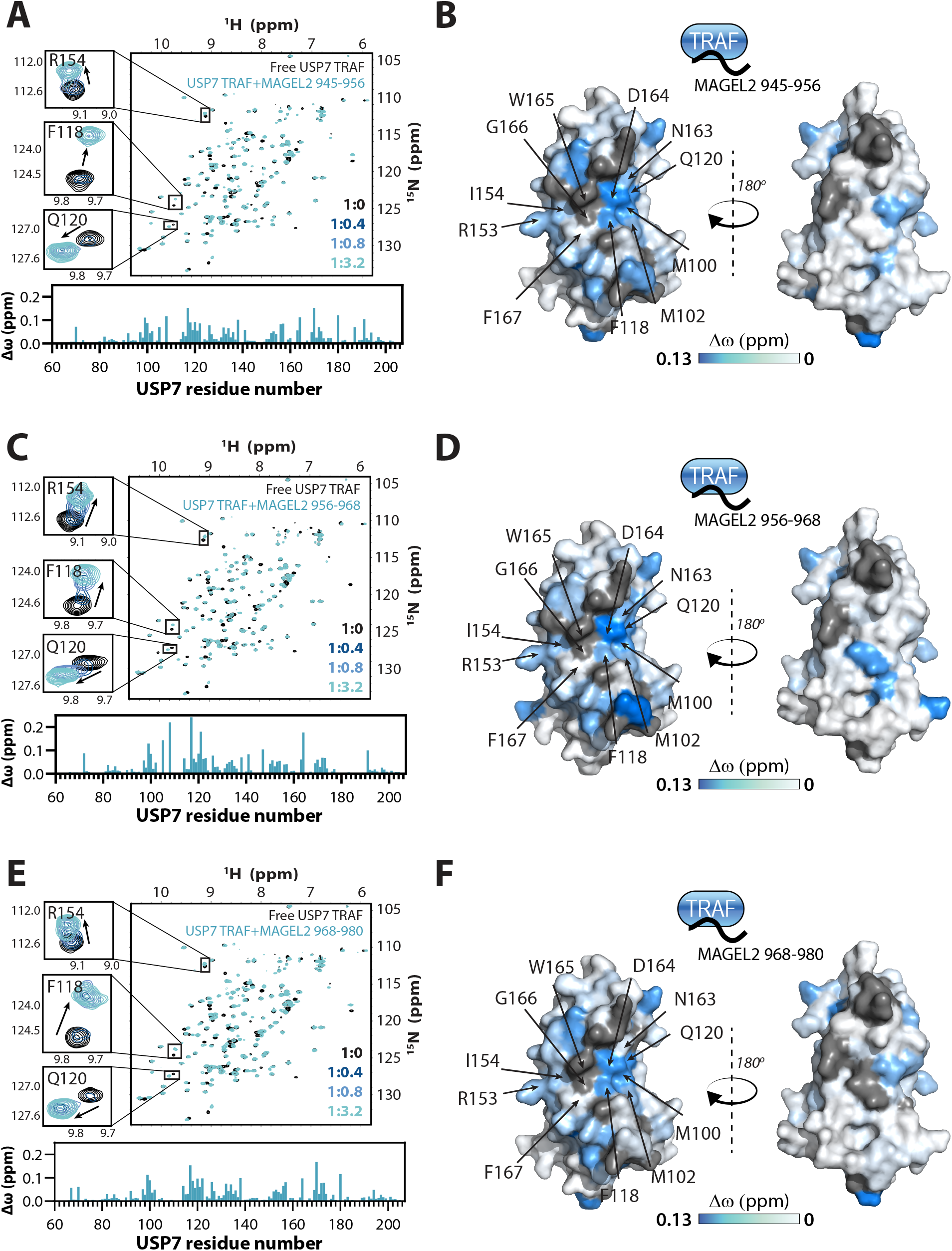
NMR mapping of MAGEL2/USP7 TRAF binding interfaces. Overlays of ^15^N-^1^H HSQC spectra of the USP7 TRAF domain alone and in complex with MAGEL2 peptides 945-968 (**A**), 956-968 (**C**), or 968-980 (**E**). Insets show peaks corresponding to TRAF residues R154, F118, and Q120 at the binding interface. USP7 to MAGEL2 molar ratios are indicated. Per-residue NMR chemical shift perturbations (Δω) are quantified as bar plots below the spectra and mapped onto the USP7 TRAF structure (PDB: 1YY6) on the right (**B, D** and **F**). The surface of the TRAF domain is color-coded according to Δω values, with more saturated blue indicating larger perturbations. Residues at the MAGEL2-binding interface are labeled.

The UBL1-2/MAGEL2 binding interface was mapped using the same approach (**Figure 4**). Addition of the unlabeled MAGEL2 peptide 882-895 caused significant perturbations in the ^15^N-^1^H TROSY spectrum of UBL1-2 (**Figure 4A-B**). The binding occurred on the slow NMR timescale, resulting in the disappearance of peaks corresponding to the free protein and the appearance of new peaks for the MAGEL2-bound state (**Figure 4A, inset**). The Δω values mapped onto the surface of UBL1-2 revealed a distinct binding interface (**Figure 4C**). It includes residues D754, K755, D758, and E759 that form the acidic patch and serve as the canonical substrate-binding site on UBL1-2^34-37,45^.

**Figure 4.**
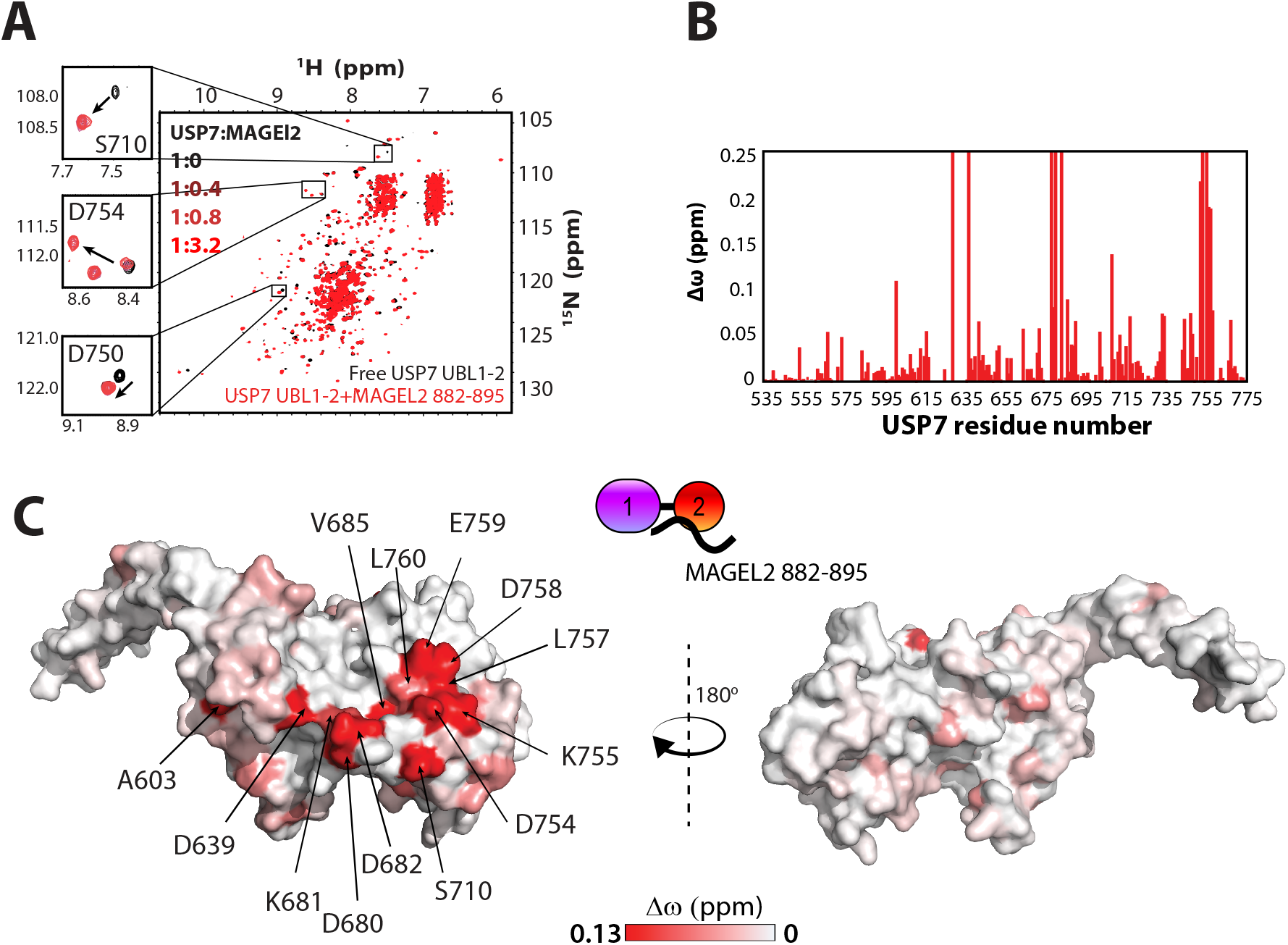
NMR mapping of MAGEL2/USP7 UBL1-2 binding interface. **A**. Overlay of ^15^N-^1^H TROSY spectra of the USP7 UBL1-2 domains alone and in complex with MAGEL2 peptide 882-895. Insets show peaks corresponding to USP7 residues S710, D754, and D750, located within the substrate binding surface of the UBL1-2 domains. USP7 to MAGEL2 molar ratios are indicated. **B**. Bar plot depicting per-residue NMR chemical shift perturbations (Δω) in the ^15^N-^1^H TROSY spectrum of the USP7 UBL1-2 domains upon the addition of a 3.2-fold molar excess of MAGEL2 882-895. Residues observed only in either “free” or “bound” states due to peak broadening were assigned to the highest observed CSP value. **C**. Mapping of Δω from (**B**) onto the USP7 UBL1-2 structure (PDB: 4PYZ) reveals the UBL1-2/MAGEL2 882-895 binding interface (red). The surface of the UBL1-2 domains is color-coded according to Δω values, with more saturated red indicating larger perturbations. Residues with high CSPs are labeled.

### Structure of the TRAF/MAGEL2 complex

To further characterize the USP7/MAGEL2 complex structure using X-ray crystallography, we attempted to crystallize the TRAF and UBL1-2 domains bound to the four chosen MAGEL2 peptides. Crystals were obtained for UBL1-2 co-crystallized with MAGEL2 fragment 882-895 and for TRAF co-crystallized with MAGEL2’s residues 968-980 (SAWEG**PSTS**RALG). While the UBL1-2 crystals did not diffract well, the TRAF crystals yielded high-quality diffraction data.

The structure of the TRAF/MAGEL2 complex was determined at a resolution of 1.63 Å with R_free_ of 0.195 and R_work_ of 0.217 (**Figure 5 and Table 1**). The TRAF domain adopts a characteristic eight-stranded antiparallel β-sandwich fold (**Figure 5A**). Clear electron density allowed MAGEL2 residues 969-AWEG**PSTS**RA-978 to be built unambiguously (**Figure 5C, right**). The peptide binds along the edge of the β-sandwich and extends the β-sheet by forming two backbone hydrogen bonds with the β7 strand of TRAF, orienting the peptide antiparallel to β7.

**Table 1.** Summary of crystallographic analysis.

| **Data Collection** | **USP7 TRAF + MAGEL2 968-980** |
| --- | --- |
| Wavelength (Å) | 0.92021 |
| Space group | *P2<sub>1</sub>* |
| Cell dimensions (Å) |  |
| *a*, *b*, *c* (Å) | 57.483, 50.696, 82.974 |
| $\alpha$, $\beta$, $\gamma$ (°) | 90.0, 103.8, 90.0 |
| Resolution (Å) | 41.499-1.628 (1.656-1.628) |
| *R*<sub>sym</sub> (%) | 0.125 (1.833) |
| Mean (*I*/σ*I*) | 6.4 (0.7) |
| Completeness (%) | 100 (100) |
| CC (1/2) | 0.995 (0.345) |
| Multiplicity | 4.3 (3.6) |
| Refinement |  |
| Resolution (Å) | 41.50-1.63 |
| No. reflections ($\vert F\vert > 0\sigma$) | 57208 |
| *R*<sub>work</sub>/*R*<sub>free</sub> (%) | 19.4/21.7 |
| No. atoms |  |
| Protein | 3,771 |
| Water | 472 |
| Average B-factors (Å<sup>2</sup>) |  |
| TRAF/MAGEL2 | 32.919/43.549 |
| Water | 39.810 |
| Wilson B-factors (Å<sup>2</sup>) | 34.23 |
| R.m.s. deviations |  |
| Bond lengths (Å) | 0.01 |
| Bond angles (°) | 1.01 |
| Ramachandran plot (%) |  |
| Favored (%) | 96.3 |
| Allowed (%) | 99.8 |
| Outliers (%) | 0.22 |

Values in parentheses are for the highest-resolution shell.

**Figure 5.**
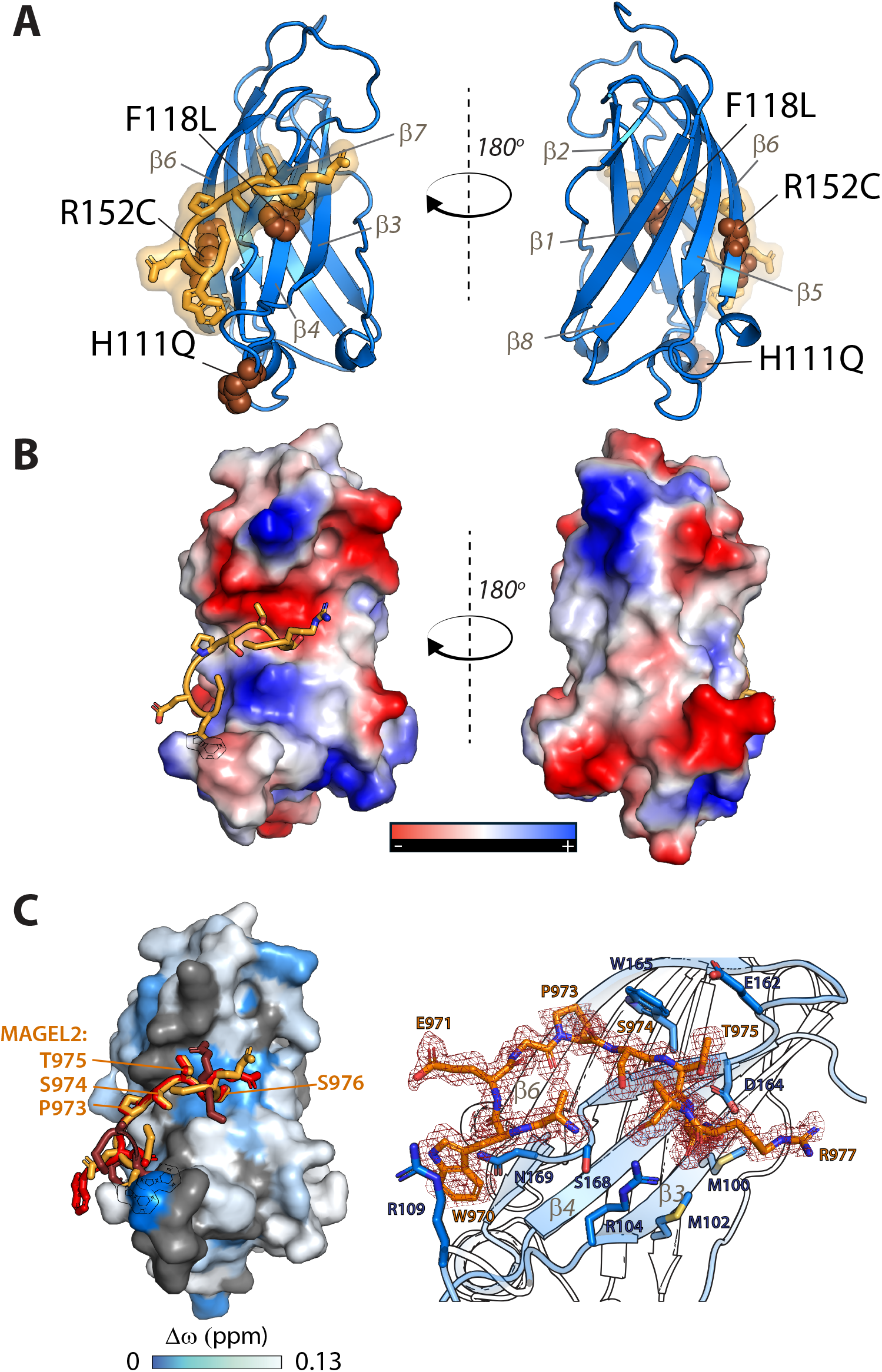
Crystal structure of MAGEL2 peptide bound to USP7 TRAF. **A**. Structure of USP7 TRAF in complex with MAGEL2 peptide 968-980 shown in ribbon representation with TRAF shown is blue and the MAGEL2 in orange. The β-strands labeled and the Hao-Fountain syndrome-linked mutations are shown as brown spheres. The isoelectric surface representation of the complex is shown in **B. C**. Structure of the TRAF surface color-coded in accordance with NMR chemical shift mapping (blue). MAGEL2 968-980 is shown as an overlay of three conformations captured in the asymmetric unit of the crystal lattice. Molecule 1/Chain B: orange; Molecule 2/Chain D: ruby; Molecule 3/Chain F: red. The PSTS motif in MAGEL2 is labeled. **D**. USP7 TRAF/MAGEL2 binding interface. The electron density of the MAGEL2 968-980 peptide is shown as dark red mesh. The interface residues of the TRAF domain making contacts with the peptide are shown as dark blue sticks. Residues of USP7 and MAGEL2 are labeled in blue and orange, respectively.

The core 973-PSTS-976 motif mediates the primary interactions. In particular, S974 forms hydrogen bonds with G166 of TRAF, anchoring the peptide to the β7 strand. The PST segment establishes an extensive network of contacts along the β7 strand, while S976 additionally interacts with residues on the β3 strand (**Figure 5C, right**). This binding mode is in good agreement with the NMR CSP mapping (**Figure 5C, left**) and consistent with previously reported crystal structures of USP7 TRAF domain bound to other substrates, confirming a conserved mechanism of substrate recognition.

The asymmetric unit contains three TRAF/MAGEL2 complexes, which exhibit differences in the conformation of residues flanking the PSTS motif; however, the core motif interactions are preserved across all models (**Figure 5C, left**). Specifically, the P973 makes extensive hydrophobic interactions with TRAF’s W165, S974 forms main chain hydrogen bonds with TRAF’s G166, T975 interacts with TRAF’s E162, and S976 forms polar interactions with the side chain of TRAF’s D164 and several close contacts with M100, M102, and R104 of TRAF. Notably, residues surrounding the PSTS motif establish additional hydrogen bonds with R104, S168, and N169 residues of TRAF. In one of the three complexes, W970 forms distinct additional interactions, including perpendicular π-stacking with Y106 and π-cation interactions with R109 of the TRAF.

These observations suggest that while the conserved PxxS motif mediates the primary interactions with the TRAF domain, flanking residues provide additional contacts that may fine-tune binding affinity and specificity, thereby contributing to selective recognition of USP7 substrates.

### The effect of disease-causing mutations on USP7/MAGEL2 interaction

After identifying the binding interfaces that mediate USP7/MAGEL2 assembly, we next tested whether Hao-Fountain syndrome-associated mutations in USP7 affect complex formation. Among reported missense variants, at least four, including H111Q, F118L, R152C, and L757P, are found in the TRAF and UBL1-2 domains (**Figure 1A**).

To evaluate their impact on MAGEL2 binding *in vitro*, we introduced individual point mutations into the isolated TRAF and UBL1-2 domains of USP7 and measured binding affinities using ITC. The MAGEL2 peptide 956-968 was used for TRAF variants, while peptide 882-895 was used for UBL1-2 variants. ITC analysis showed that the H111Q and R152C mutations do not significantly affect the binding, with affinities comparable to the WT (4.29±1.38 μM and 5.60±3.26 μM, respectively). In contrast, the F118L mutation results in a dramatic decrease in binding (**Figure 6A**). Similarly, the L757P mutation in the UBL1-2 domain completely abolished binding to MAGEL2 (**Figure 6B**).

**Figure 6.**
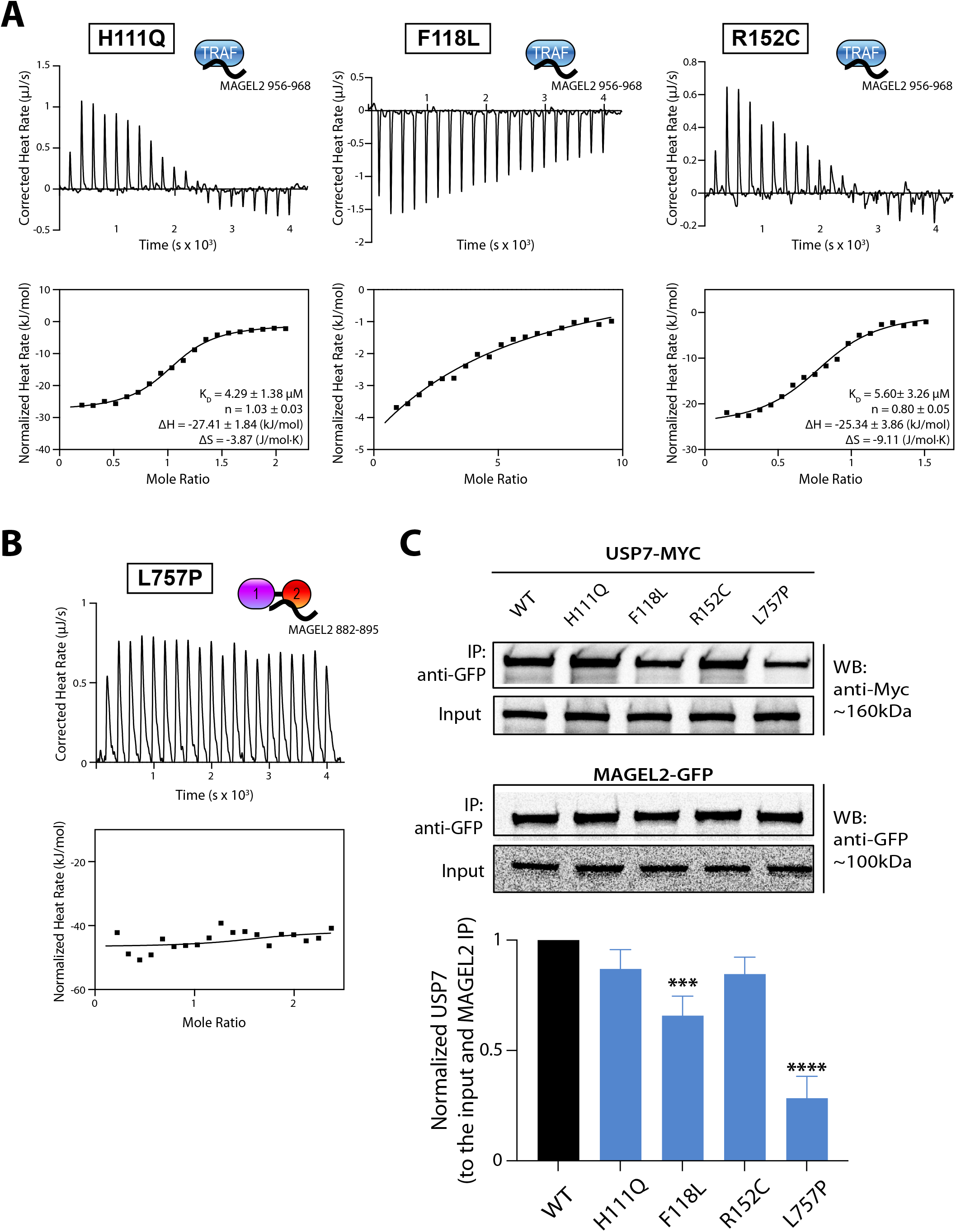
Hao-Fountain syndrome-linked mutations affect USP7/MAGEL2 complex formation. **A**. ITC binding isotherms of the USP7 TRAF and its H111Q, R152C, and F118L mutants titrated with MAGEL2 peptide 956-968. **B**. ITC binding isotherm of L757P mutant in UBL1-2 titrated with MAGEL2 peptide 882-895. **C**. Co-immunoprecipitation of MAGEL2 with USP7 and its Hao-Fountain syndrome-linked variants in HEK293FT cells. GFP-tagged MAGEL2 was co-expressed in 293FT cells with WT or mutant Myc-tagged USP7. Cell lysates were subjected to co-immunoprecipitation using anti-GFP antibody, and immunoprecipitated complexes were analyzed by Western blotting with anti-Myc and anti-GFP antibodies. Input and immunoprecipitated samples are shown. Quantification of USP7 binding (normalized to input and immunoprecipitated MAGEL2) is shown relative to WT (set to 1). Statistical significance is indicated (*** p-value<0.001, **** p-value<0.0001). Data represent three independent experiments for all.

These interactions were further validated by orthogonal NMR titration experiments. Consistent with the ITC results, NMR revealed that the H111Q and R152C mutations exhibited pronounced chemical shift perturbations, comparable to the WT, indicative of TRAF binding. In contrast, the F118L mutation displayed markedly fewer and smaller CSPs, suggesting reduced binding (**Supplementary Figure S2**). Comparison of the free ^15^N-^1^H HSQC spectra of the WT and mutant TRAF domains showed that the H111Q, F118L, and R152C mutations caused minimal perturbations to the NMR spectra, indicating that these substitutions did not compromise the overall structural integrity of the domain (**Supplementary Figure S2**). In contrast, ^15^N-^1^H TROSY spectrum for the UBL1-2 mutant revealed poor spectral quality compared to the WT protein, indicative of major folding defects that are detrimental to UBL1-2’s ability to bind MAGEL2 (**Supplementary Figure S3**).

To assess these effects in a cellular context, we performed co-immunoprecipitation (co-IP) assays. HEK293FT cells were co-transfected with GFP-tagged MAGEL2 and Myc-tagged USP7 variants, followed by co-immunoprecipitation with anti-GFP antibody (**Figure 6C)**. Consistent with the *in vitro* finding, H111Q and R152C variants show little to no effect on USP7/MAGEL2 interaction, whereas F118L significantly reduces binding, and the L757P mutation in the UBL1-2 domain nearly abolishes complex formation. Overall, the cellular experiments complement our biophysical findings.

Together, these data demonstrate that disease-associated mutations in USP7 can compromise USP7 function by impairing MAGEL2 complex formation.

## DISCUSSION

Mutations and deletions in MAGEL2 are associated with Prader-Willi and Schaaf-Yang syndromes^14-17^. More recently, mutations in USP7 have been linked to Hao-Fountain syndrome, which presents with significant phenotypic overlap with the former two conditions^11,20^. While Prader-Willi and Schaaf-Yang syndromes have been extensively studied, HAFOUS remains poorly understood. Building on the findings of Hao et al.^11^, who identified key interactions within the MUST complex, we aimed to characterize the specific binding mechanism of MAGEL2 recognition by USP7 and to evaluate how disease-relevant USP7 variants may affect this interaction.

This work demonstrates that USP7 interacts with MAGEL2 in a bipartite manner. We identified two distinct linear USP7-binding motifs within the intrinsically disordered N-terminal region of MAGEL2, each recognized by a different USP7 domain. The conserved KATRKKK motif within residues 882-895 is bound by the UBL1-2 domains, while the TRAF domain exhibits broader specificity, engaging six of the seven E/PxxS motifs in this region. Notably, a distinct section of MAGEL2 encompassing residues 945-980 is highly conserved and includes three TRAF-binding peptides (945-956, 956-968, and 968-980), underscoring their likely functional significance. Interestingly, these three peptides share similar sequences beyond the canonical consensus, each featuring a tryptophan, glutamic acid, and glycine flanking the N-terminal region, which likely accounts for their comparable binding affinities (**Figure 1B**). The ability for TRAF to engage multiple motifs, together with the similar affinities for each of the three studied TRAF-binding peptides, supports a dynamic multivalent binding model in which the TRAF domain of USP7 engages multiple proximal motifs within MAGEL2, potentially enhancing overall avidity and specificity.

Given the central role of the MUST complex in neurodevelopment, we hypothesized that disruption of the USP7/MAGEL2 interaction may contribute to the shared phenotypic features of USP7-related Hao-Fountain syndrome and MAGEL2-related Prader-Willi and Schaaf-Yang syndromes. Using NMR and ITC, we demonstrate that the Hao-Fountain syndrome-associated F118L mutation in the TRAF domain of USP7 nearly abolishes MAGEL2 binding *in vitro* without affecting overall protein folding or stability (**Figure 6A** and **Supplementary Figure S2**). This is well supported by the crystal structure of TRAF in complex with the MAGEL2 peptide (**Figure 5A**), which shows that F118 lies directly on the MAGEL2 binding interface. In contrast, the H111Q and R152C mutations located adjacent to the binding interface captured in the crystal structure have only a marginal negative effect on binding. In good agreement with the *in vitro* results, F118L USP7 has decreased interaction with MAGEL2, while H111Q and R152C have no significant effect on MAGEL2 binding in cells.

Unlike the TRAF variants tested here, the L757P mutation in the isolated UBL1-2 domains induces dramatic changes in the NMR spectra, consistent with protein aggregation likely due to folding defects caused by the leucine-to-proline substitution at the UBL1/UBL2 inter-domain interface. Consistent with this, ITC and co-IP data show that the L757P variant of USP7 abolishes binding to MAGEL2 both *in vitro* and in cells (**Figure 6B-C** and **Supplementary Figure S3**).

The bipartite mode of interaction between MAGEL2 and USP7 implies that impairment of one binding site may be at least partially compensated by the other. Indeed, although the F118L mutation in the isolated TRAF domain nearly abolishes binding to a MAGEL2-derived peptide *in vitro*, it results in only a 30-50% reduction in co-immunoprecipitation in cells, suggesting that the UBL1-2 domains can partially compensate for reduced TRAF-mediated binding. On the other hand, the binding-deficient L757P UBL1-2 variant has a more pronounced effect on USP7/MAGEL2 complex formation, causing an ∼70% reduction in co-immunoprecipitation. This further highlights the importance of the conserved KxxxKxK motif in MAGEL2 for stable complex formation (**Figure 6C**).

Schaaf-Yang syndrome is caused by truncating mutations in MAGEL2, most of which occur within its C-terminal half (residues ∼500-1249)^16,17,46,47^. As we have shown, MAGEL2 engages USP7 in a bipartite manner, with both the TRAF and UBL1-2 binding regions spanning residues 882-1008 within the USP7-BR region. Notably, these sites fall within the region affected by Schaaf-Yang-associated truncations. Loss of either binding site is expected to disrupt USP7 interaction. This shared mechanism of MUST complex disruption offers a plausible molecular explanation for the clinical similarities between Schaaf-Yang and Hao-Fountain syndromes. Finally, while some USP7 variants analyzed here do not significantly impair MAGEL2 binding, they may affect other interactions within the MUST complex, not tested in this study, such as regions outside out the USP7-BR and TRIM27. Therefore, pathogenicity may arise from disruptions at multiple points within the complex.

Overall, this work represents an important first step in understanding the molecular architecture of the MUST complex. By defining how USP7 and MAGEL2 interact and how disease-linked mutations disrupt this interaction, we provide a mechanistic basis for the clinical overlap between Hao-Fountain and Schaaf-Yang syndromes. These findings lay the foundation for future studies on how mutations in other components of the MUST complex contribute to neurodevelopmental disorders. Understanding the molecular basis of disorders like Hao-Fountain, Schaaf-Yang, and Prader-Willi syndromes improves diagnosis and opens potential therapeutic avenues.

## RESOURCE AVAILABILITY

The atomic coordinates and structure factors of USP7 TRAF in complex with MAGEL2 (968-980) have been deposited in the Protein Data Bank (http://www.wwpdb.org/) with PDB ID 12ZJ. All plasmids for bacterial expression of USP7 variants were deposited to Addgene database. 227226 (WT USP7 TRAF), 227227 (H111Q USP7 TRAF), 227228 (F118L USP7 TRAF), 227229 (R152C USP7 TRAF), 227230 (WT USP7 UBL1-2), 227231 (L757P USP7 UBL1-2). Mammalian expression plasmids are available upon request.

## ACKNOWLEDGEMENTS

We would like to thank Dr. Vivian Saridakis for providing us the UBL1-2 construct and Dr. Dmitry Korzhnev for NMR support.

## AUTHOR CONTRIBUTIONS

E.J.K. and I.B. wrote the manuscript and prepared figures. E.J.K. designed and generated all USP7 mutants used for NMR and ITC. E.J.K. designed and performed all NMR assays. E.J.K. crystallized USP7 TRAF in complex with MAGEL2. E.J.K. and B.H. refined the crystal structure. E.J.K. and G.A.S. designed and performed all ITC assays. I.S. designed and generated all cellular USP7 mutants. D.S. and T.B. performed the cellular assays. I.B. and K.F.T. conceived the cellular work. I.B. conceptualized the project, acquired funding, reviewed, and edited the manuscript. All authors read and approved the final version prior to submission.

## MATERIALS AND METHODS

### Protein expression and purification

Plasmids for cellular expression of USP7 contained the full-length USP7 coding sequence inserted into a pCS2+ vector downstream of a Myc tag, while the MAGEL2 sequence encoding the last 645 amino acids^9,11^ was inserted into a pIRESpuro downstream of a GFP tag. Plasmids for bacterial expression of USP7 and its mutants contained either the coding sequence of the TRAF domain of USP7 (62-205) or the UBL1-2 domains (535-775). The TRAF domain was inserted into a pET28b+ vector (Genscript), downstream of the N-terminal 6xHis tag and thrombin and TEV cleavage sites. The UBL1-2 domains were inserted into a pET15b vector (Vivian Saridakis), downstream of the N-terminal 6xHis tag and TEV cleavage site. Hao-Fountain syndrome mutations were introduced using site-directed mutagenesis. The constructs were sequence-verified and the plasmids for bacterial expression were deposited in the Addgene database.

USP7 constructs were transformed into Escherichia Coli BL21 (DE3) cells and expressed in 1L Luria Broth (LB) media for ITC. For NMR titration experiments, ^15^N-labeled proteins were expressed in M9 minimal media with ^15^NH_4_Cl as the sole nitrogen source. Cells were grown at 37°C until OD_600_ reached 0.8-1.0 o.u. Protein expression was induced with 1 mM IPTG at 20°C overnight. Cells were harvested by centrifugation, resuspended in lysis buffer (20 mM sodium phosphate pH 8.0, 250 mM NaCl, 5 mM imidazole, 0.5 mM PMSF), and lysed by sonication. The lysate was centrifuged at 15,000 rpm for 45 minutes, and the supernatant was filtered and applied to TALON HisPur cobalt resin (Thermo Scientific). USP7 constructs were eluted with a buffer containing 20 mM sodium phosphate pH 8.0, 250 mM NaCl, and 250 mM imidazole. TEV was added to remove the 6-His tag, and the samples were buffer-exchanged overnight at 4°C into 20 mM sodium phosphate pH 7.4, and 250 mM NaCl to remove imidazole and allow for cleavage. Proteins were then purified by size-exclusion chromatography using a HiLoad Superdex 75 column (GE Healthcare) and eluted in 20 mM phosphate pH 8.0, and 250 mM NaCl for USP7 TRAF domains and 20 mM phosphate pH 7.0, and 100 mM NaCl for UBL1-2 domains.

MAGEL2 peptides were purchased from Genscript and resuspended in appropriate buffers corresponding to either TRAF or UBL1-2.

### NMR titration experiments

NMR data were collected on a Bruker Avance NEO 600 and 800 MHz equipped with a cryoprobe at 30°C for USP7 TRAF (600 MHz) and 25°C for UBL1-2 (800 MHz). Data processing was performed with NMRPipe^48^, and analysis was done using Sparky^49^. The USP7 TRAF domains were in a buffer containing 20 mM phosphate pH 8.0, 250 mM NaCl, 10 mM DTT, and 10% D2O (v/v). The USP7 UBL1-2 domains were in a buffer containing 20 mM phosphate pH 7.0, 100 mM NaCl, 10 mM DTT, and 10% D2O (v/v).

To study MAGEL2 peptides binding, unlabeled peptides were gradually added to ∼150-250 µM ^15^N-labeled USP7 up to 1:3.2 molar excess of peptide. Chemical shift changes were monitored using 2D ^15^N-^1^H HSQC (TRAF) or TROSY (UBL1-2) spectra across seven titration points. Chemical shift perturbations were plotted using GraphPad Prism and mapped on the structures of TRAF and UBL1-2 using PyMOL.

### Isothermal Titration Calorimetry (ITC)

All measurements were collected using an Affinity ITC instrument (TA Instruments). ITC data were analyzed with NanoAnalyze software (TA Instruments). Data fitting was performed using an “independent” model after correcting for the heat of peptide’s dilution. USP7 TRAF was in a buffer containing 20 mM phosphate buffer pH 8.0 and 250 mM NaCl. MAGEL2 peptides containing the A/PxxS motif were resuspended in this same buffer and titrated into ∼100 µM USP7 TRAF WT and mutants. Experiments were performed at 4°C with a total of 20 injections made with 200-second time intervals and 200 rpm mixing speed. USP7 UBL1-2 was in a buffer containing 20 mM phosphate buffer pH 7.0 and 100 mM NaCl. The MAGEL2 peptide, containing KxxxKxK motif, was resuspended in the same buffer and titrated into ∼50µM USP7 UBL1-2 WT and L757P mutant. A total of 20 injections were made with 200 second time intervals and 200 rpm mixing speed at 25°C.

### X-ray crystallography

Seed crystals of USP7 TRAF (22 mg/mL) in complex with MAGEL2 968-980 peptide (1:2 ratio protein to peptide) were first obtained in 250 mM MgCl_2_, 100 mM Tris pH 9.0, and 30%PEG 8000 at 16 °C using hanging-drop vapor diffusion and then crushed through shaking with small steel beads using a vortex. Larger USP7 TRAF crystals (22 mg/mL) in complex with MAGEL2 968-980 (1:2 ratio protein to peptide) were then obtained in 250 mM MgCl_2_, 100 mM Tris pH 9.0, and 30%PEG 8000 at 16°C with seeding through hanging-drop vapor diffusion. Crystals were cryoprotected in reservoir solution supplemented with 20% glycerol and flash-cooled in liquid nitrogen. X-ray diffraction data were collected at NSLS-II beamlines 17-ID-1 and 17-ID-2 and processed using Fast DP^50^ and autoPROC^51^. The crystals contain three molecules in the asymmetric unit. The structure of the USP7 TRAF was determined by molecular replacement in Phaser/CCP4^52-54^ using the published USP7 TRAF structure (PDB ID 2FOJ)^42^ as a starting model and the resulting electron density map was readily interpretable to build in the MAGEL2 peptide. The models were refined by alternating cycles of manual rebuilding in Coot and refinement with REFMAC5/CCP4^52-57^. Data collection and refinement statistics are summarized in **Table 1**. Ramachandran statistics were calculated using MolProbity^58^, and figures generated in PyMOL^59^ (Schrödinger, LLC).

### Cell culture and transfection

HEK293FT cells were cultured in 6-well culture plates (Corning, 07-200-83) in high glucose Dulbecco’s Modified Eagle Medium (Gibco, 11065092) supplemented with 10% regular fetal bovine serum (Corning, 335-010-CV), 100 units/ml penicillin (Gibco, 15140-122) and 100 µg/ml streptomycin (Gibco, 15140-122). Plasmid transfection was performed using Lipofectamine™ 3000 transfection reagent (ThermoFisher Scientific, L3000015) according to the manufacturer’s instructions.

### Cell lysis, Co-Immunoprecipitation, and Western blot analysis

Cells were collected and lysed in Pierce IP Lysis Buffer (ThermoFisher Scientific, 87788) by incubation on ice for 20 minutes, followed by end-over-end mixing at 4°C for 40 minutes. A 30 µl aliquot of each lysate was mixed with 10 µl of 4X Laemmli Sample Buffer (Bio rad, 1610747) and used as input control. 450 µl of each sample was incubated with 2.5 µl of anti-GFP antibody (ab6556, Abcam) for 3 h at 4°C with end-over-end mixing. 50 µl of Pierce Protein A agarose (ThermoFisher Scientific, 20333) was added to each sample and incubated for 1 hour at 4°C with end-over-end mixing.

Protein complexes were eluted by boiling the agarose beads in 40 µl of 2X Laemmli Sample Buffer (Millipore Sigma, S3401) at 95°C for 5 min. The proteins were separated on 10% polyacrylamide gels (Bio rad, 4561033) at 80 V and transferred to 0.2 µm nitrocellulose membranes using the Trans-Blot Turbo Transfer System (Bio rad, 1704150) and Trans-Blot Turbo Transfer Packs (Bio rad, 1704158).

Membranes were blocked with 5% BSA in TBST for 1 h at room temperature and incubated overnight with anti-MYC antibody (homemade, 0.8 mg/ml) diluted 1:1000 in 5% BSA/TBST, at 4°C with gentle rocking. After washing, the membranes were incubated with anti-mouse IgG HRP-linked secondary antibody (Cell Signaling Technology, 7076S) diluted 1: 10 000 in 5% BSA/TBST, at room temperature for 1 hour. Protein bands were detected using ECL Western Blotting Detection Reagents (Cytiva, RPN2209) and imaged on the Azure 600 Western Blot Imager.

Bands were quantified using ImageJ (v2.16.0/1/54p). The intensity of USP7-myc immunoprecipitated bands was normalized to the corresponding bands in the 2% input and to the immunoprecipitated MAGEL2-GFP band. USP7 WT was set to 1. Statistical analyses were conducted using a one-way ANOVA followed by Dunnett’s multiple comparison test on GraphPad Prism.

## ABBREVIATIONS

DUB: deubiquitinating enzyme
USP: ubiquitin-specific protease.
TRAF: tumor necrosis factor receptor-associated factor
UBL: ubiquitin-like
WASH complex: Wiskott Aldrich Syndrome protein and scar homologue complex
MAGEL2: MAGE family member L2
TRIM27: Tripartite Motif Containing 27
HAFOUS: Hao-Fountain syndrome
NMR: Nuclear Magnetic Resonance
ITC: Isothermal Titration Calorimetry
HSQC: Heteronuclear Single Quantum Coherence
TROSY: Transverse Relaxation Optimized Spectrometry
CSP: Chemical Shift Perturbations
K_D_: Dissociation constant
WT: wild type
TEV: Tobacco Etch Virus Protease
Co-IP: Co-immunoprecipitation

