## Supplemental Data for "Mechanisms of USP7/MAGEL2 Complex Assembly and Its Mutational Disruption in Neurodevelopmental Diseases"

### Figure S1

**A**

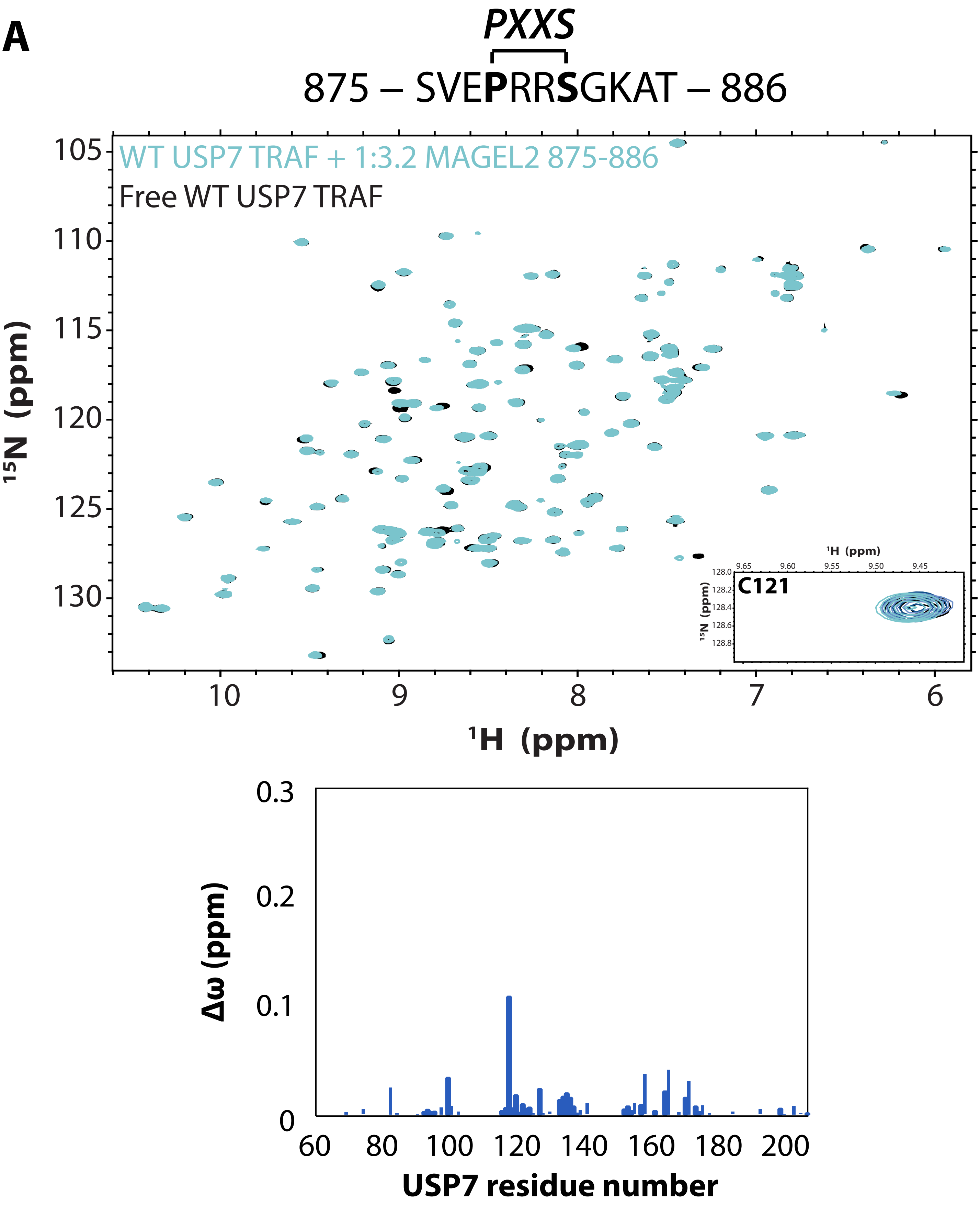
PXXS
875 – SVEPRRSGKAT – 886
WT USP7 TRAF + 1:3.2 MAGEL2 875-886
Free WT USP7 TRAF
C121
$^{15}$N (ppm)
$^{1}$H (ppm)
Δω (ppm)
USP7 residue number

Figure S1

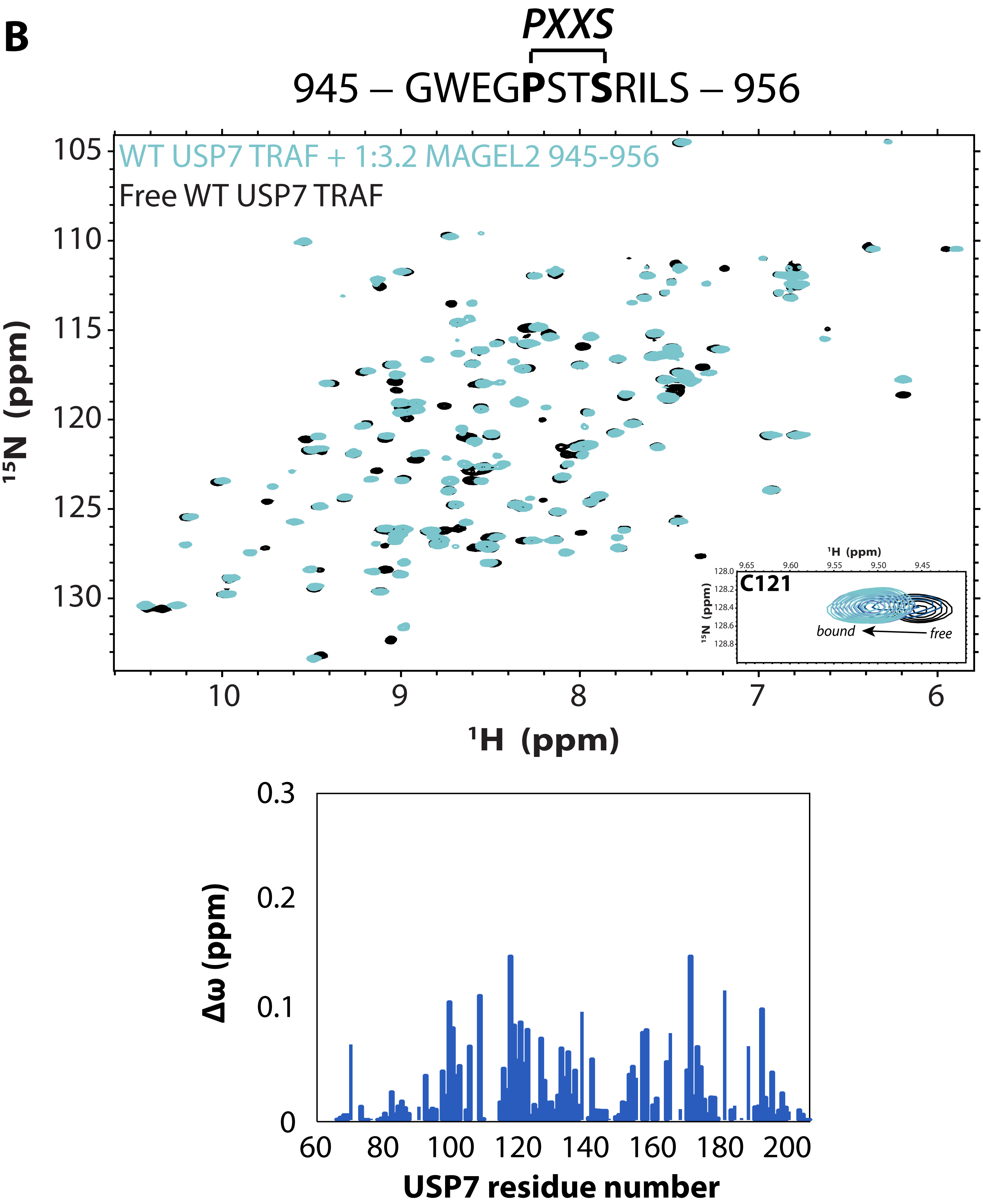
B
PXXS
945 – GWEGPSTSRILS – 956
WT USP7 TRAF + 1:3.2 MAGEL2 945-956
Free WT USP7 TRAF
15N (ppm)
1H (ppm)
C121
bound
free
Δω (ppm)
USP7 residue number

Figure S1

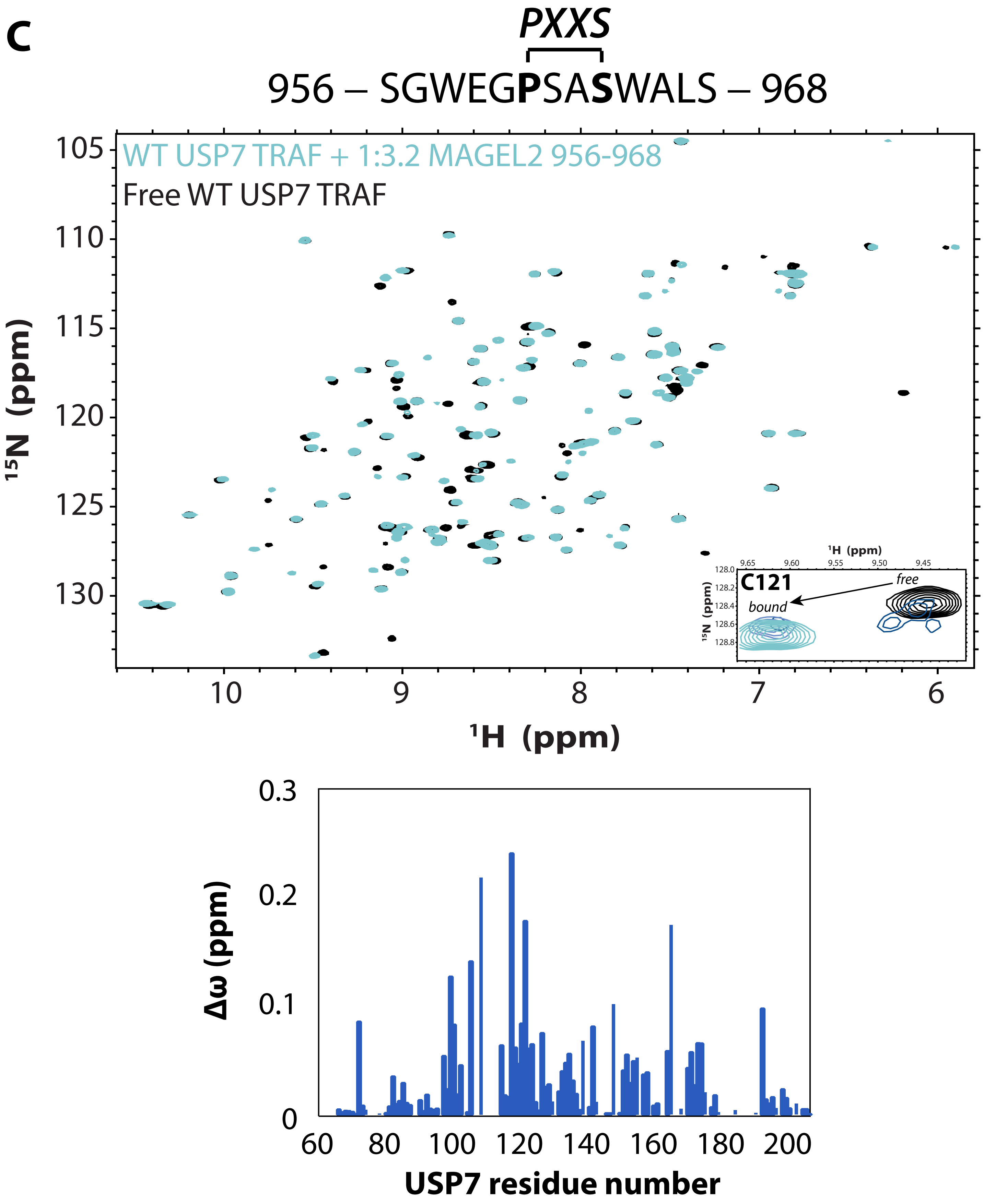
C
PXXS
956 – SGWEGPSASWALS – 968
WT USP7 TRAF + 1:3.2 MAGEL2 956-968
Free WT USP7 TRAF
15N (ppm)
1H (ppm)
C121
free
bound
Δω (ppm)
USP7 residue number

Figure S1

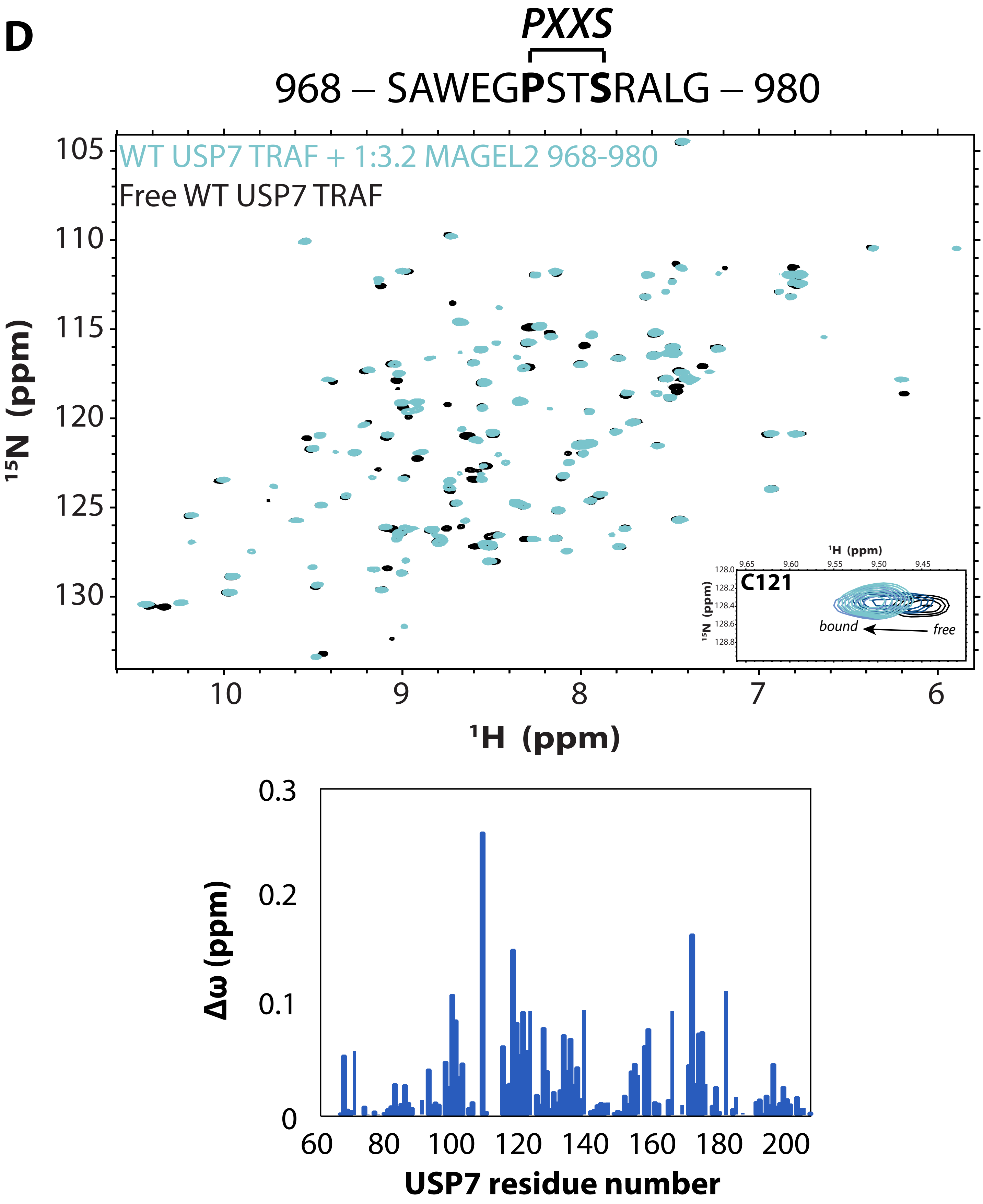
D
PXXS
968 – SAWEGPSTSRALG – 980
WT USP7 TRAF + 1:3.2 MAGEL2 968-980
Free WT USP7 TRAF
105
110
115
120
125
130
15N (ppm)
10
9
8
7
6
1H (ppm)
C121
bound
free
1H (ppm)
15N (ppm)
0.3
0.2
0.1
0
Δω (ppm)
60
80
100
120
140
160
180
200
USP7 residue number

### Figure S1

**E**

*PXXS*

982 – SES**P**GS**S**LPVVVSE – 995

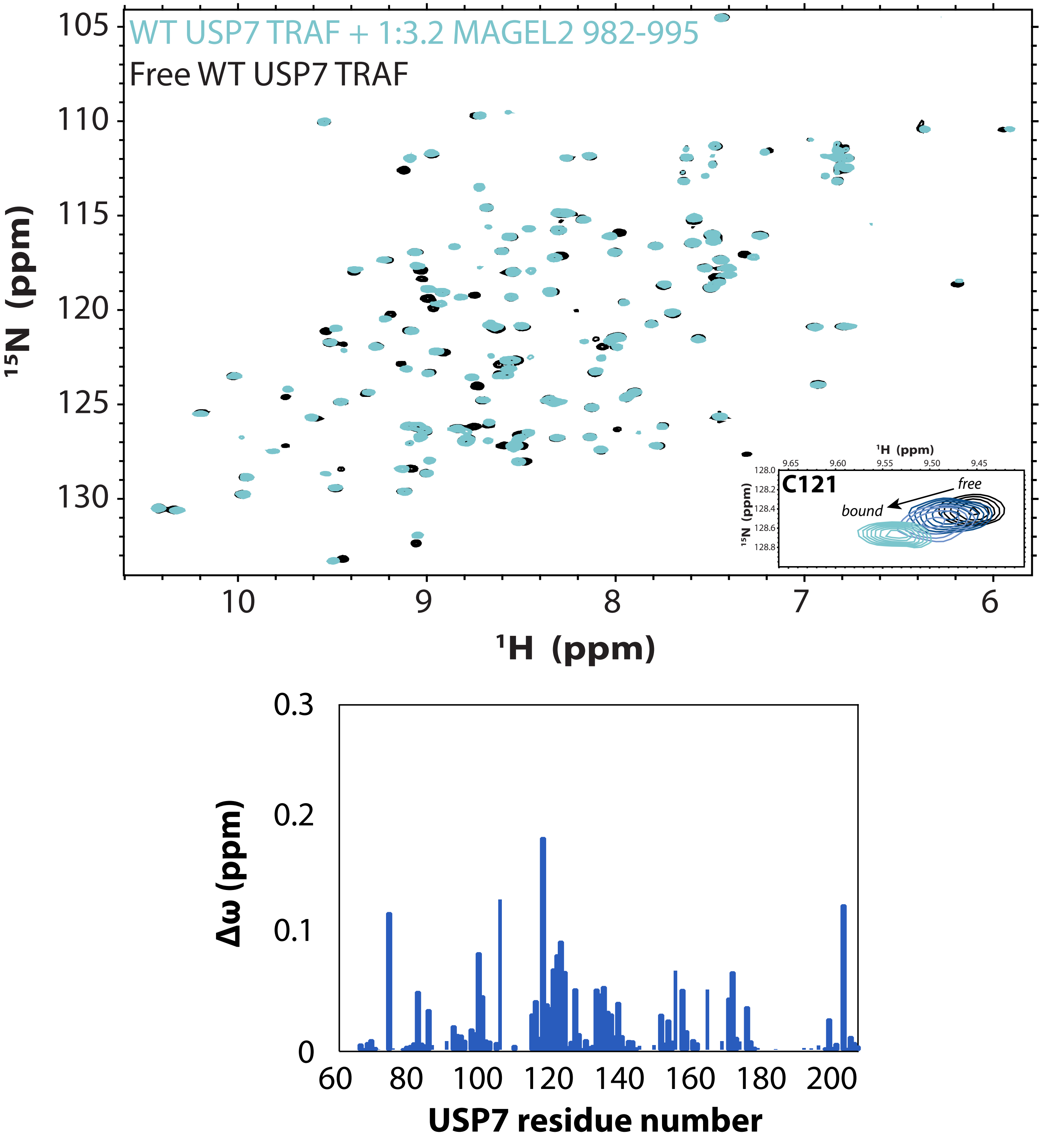

Figure S1

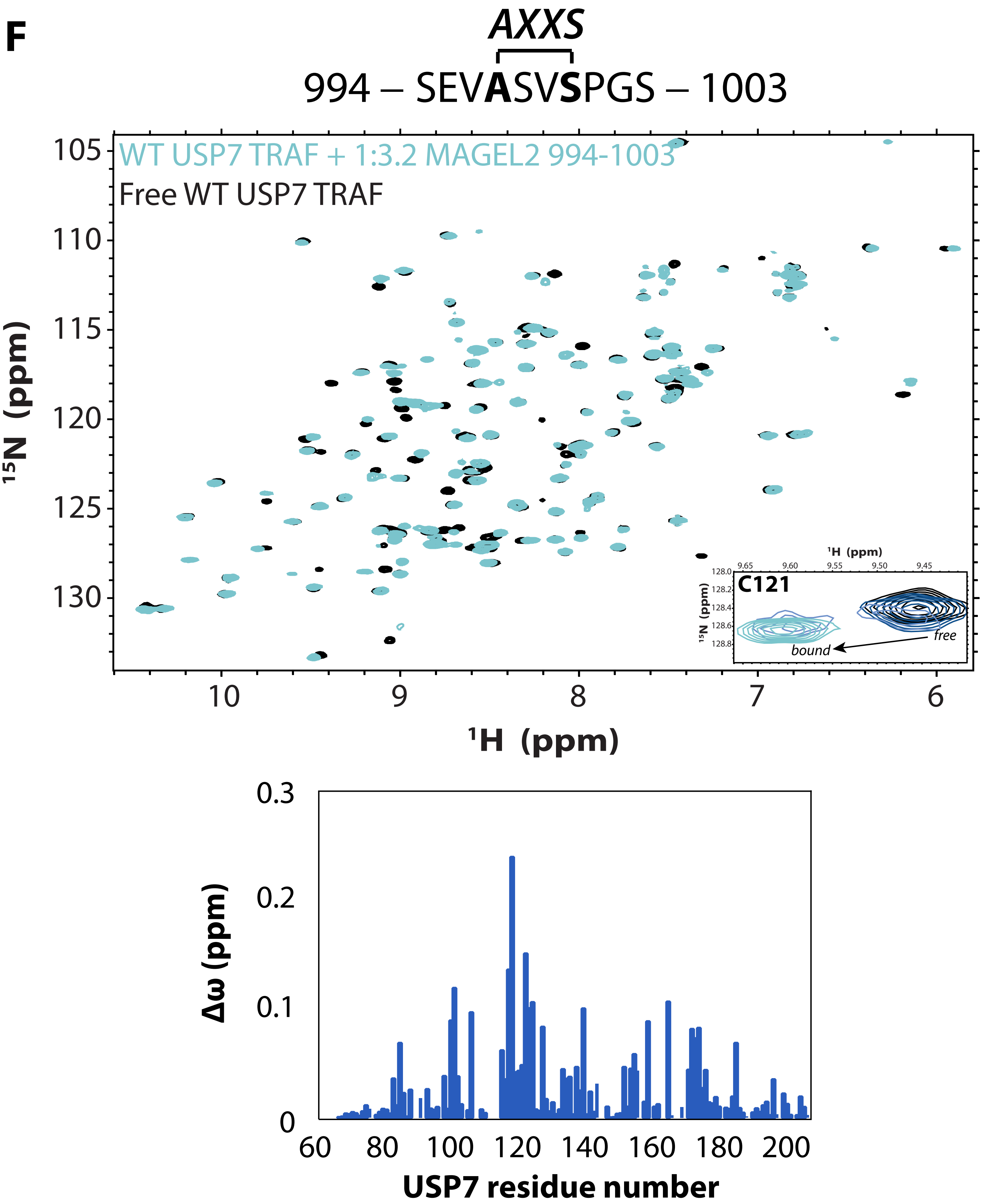
F
AXXS
994 – SEVASVSPGS – 1003
WT USP7 TRAF + 1:3.2 MAGEL2 994-1003
Free WT USP7 TRAF
C121
bound
free
¹H (ppm)
¹⁵N (ppm)
Δω (ppm)
USP7 residue number

### Figure S1

**G**

*PXXS*

998 – SVS**P**GS**S**ATQD – 1008

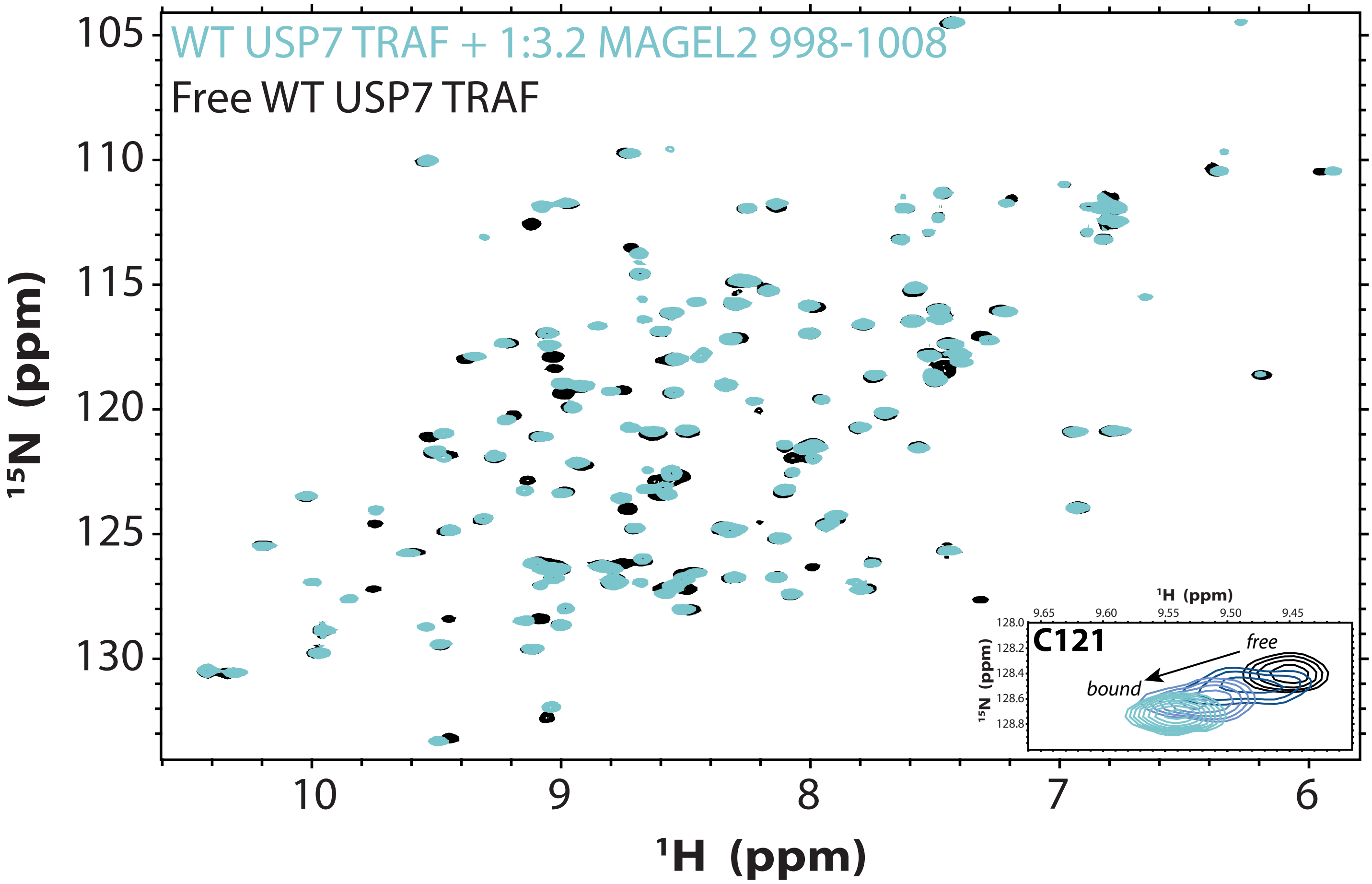


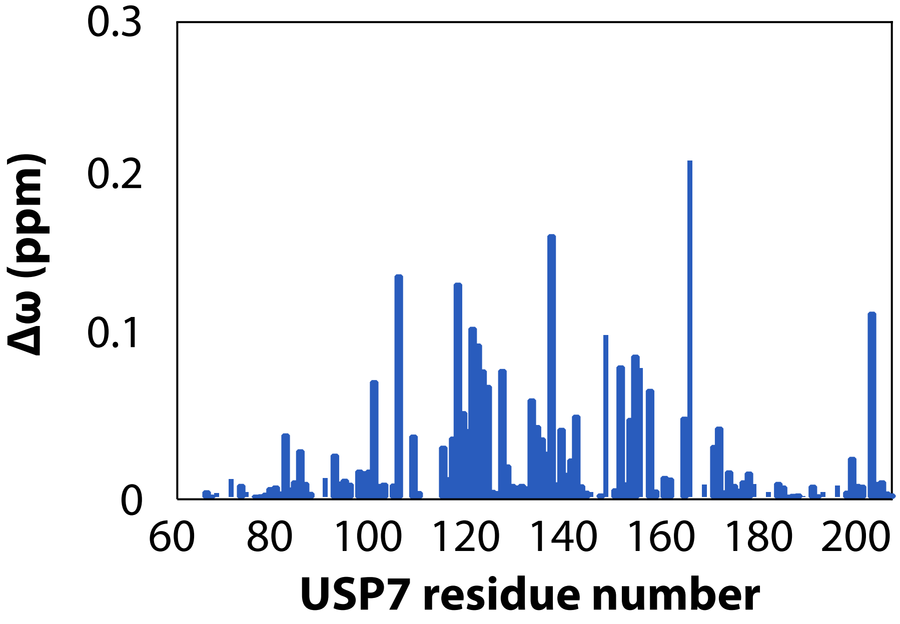

Figure S1

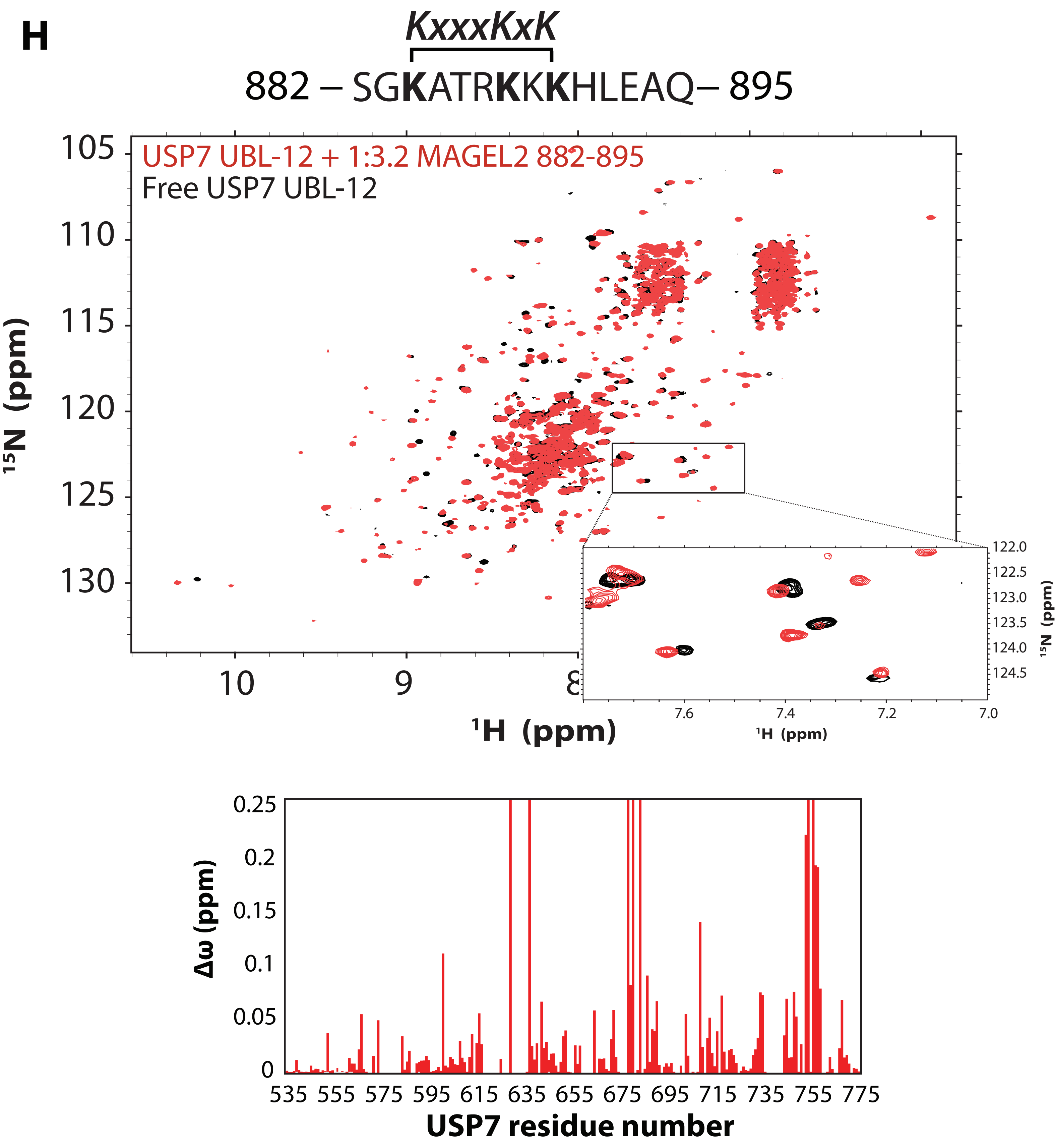
H
KxxxKxK
882 – SGKATRKKKHLEAQ– 895
USP7 UBL-12 + 1:3.2 MAGEL2 882-895
Free USP7 UBL-12
15N (ppm)
1H (ppm)
105
110
115
120
125
130
10
9
8
122.0
122.5
123.0
123.5
124.0
124.5
7.6
7.4
7.2
7.0
Δω (ppm)
0.25
0.2
0.15
0.1
0.05
0
535 555 575 595 615 635 655 675 695 715 735 755 775
USP7 residue number

**Figure S1. NMR analysis of interactions between USP7 and peptides derived from the USP7-binding region of MAGEL2.**

Overlays of $^{15}N$-$^{1}H$ HSQC spectra of the USP7 TRAF domain alone (black) and in the presence of MAGEL2 peptides (blue) at a 3.2 molar excess of peptides 875-886 (***A***), 945-956 (***B***), 956-968 (***C***), 968-980 (***D***), 982-995 (***E***), 994-1003 (***F***), and 998-1008 (***G***). Insets show a representative peak corresponding to C121. Per-residue NMR chemical shift perturbations (Δω) are quantified and shown as bar plots below each spectral overlay. **H.** Overlay of $^{15}N$-$^{1}H$ HSQC spectra of the USP7 UBL1-2 domains alone (black) and in the presence of a 3.2 molar excess of MAGEL2 peptide 882-895 *(*red*)*. The corresponding per-residue Δω values are shown as a bar plot below the spectral overlay.

### Figure S2

**A**

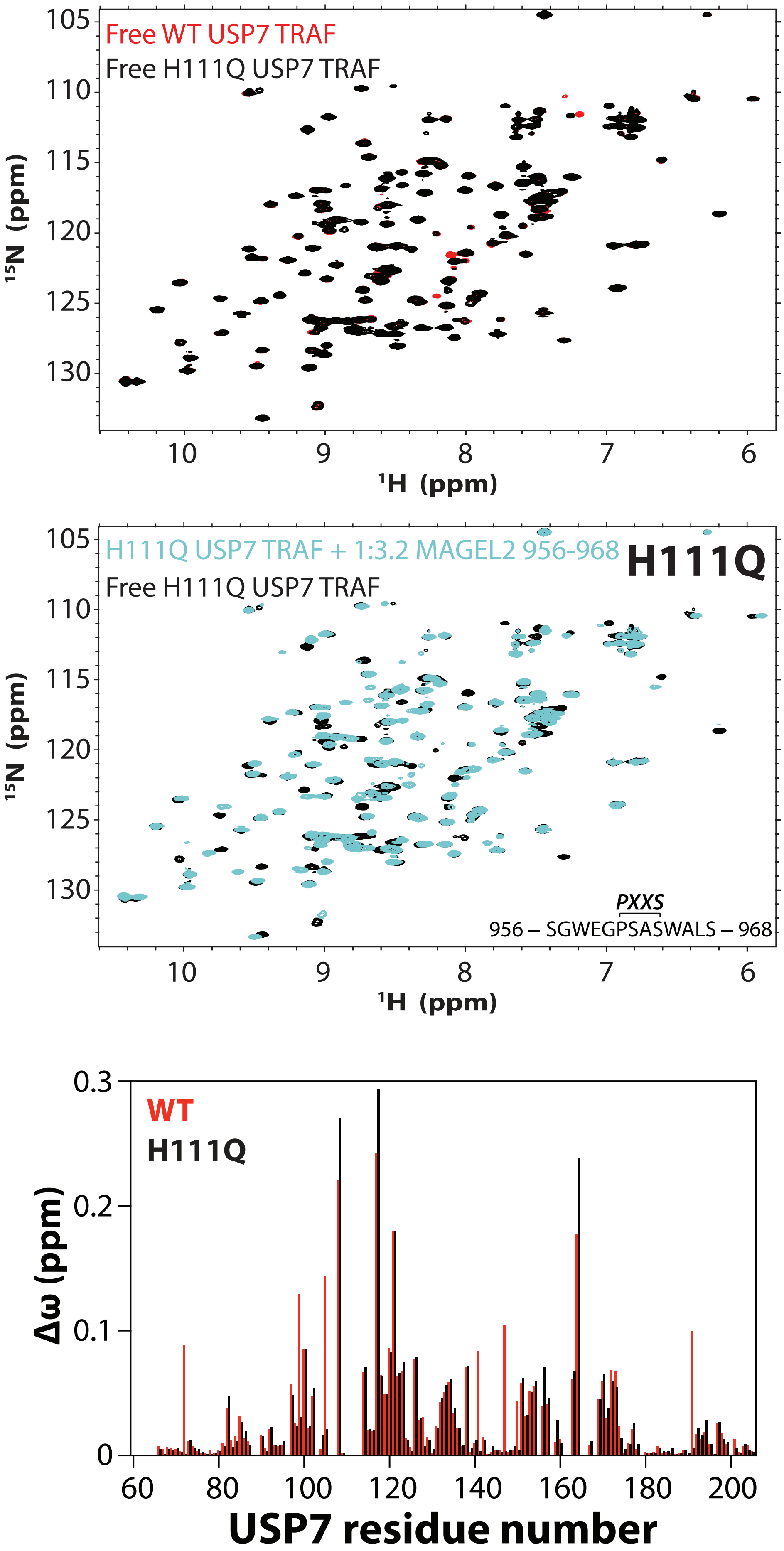

Free WT USP7 TRAF
Free H111Q USP7 TRAF
$^{15}$N (ppm)
$^{1}$H (ppm)
105
110
115
120
125
130
10
9
8
7
6
H111Q USP7 TRAF + 1:3.2 MAGEL2 956-968
Free H111Q USP7 TRAF
H111Q
PXXS
956 – SGWEGPSASWALS – 968
WT
H111Q
Δω (ppm)
0.3
0.2
0.1
0
60
80
100
120
140
160
180
200
USP7 residue number

### Figure S2

**B**

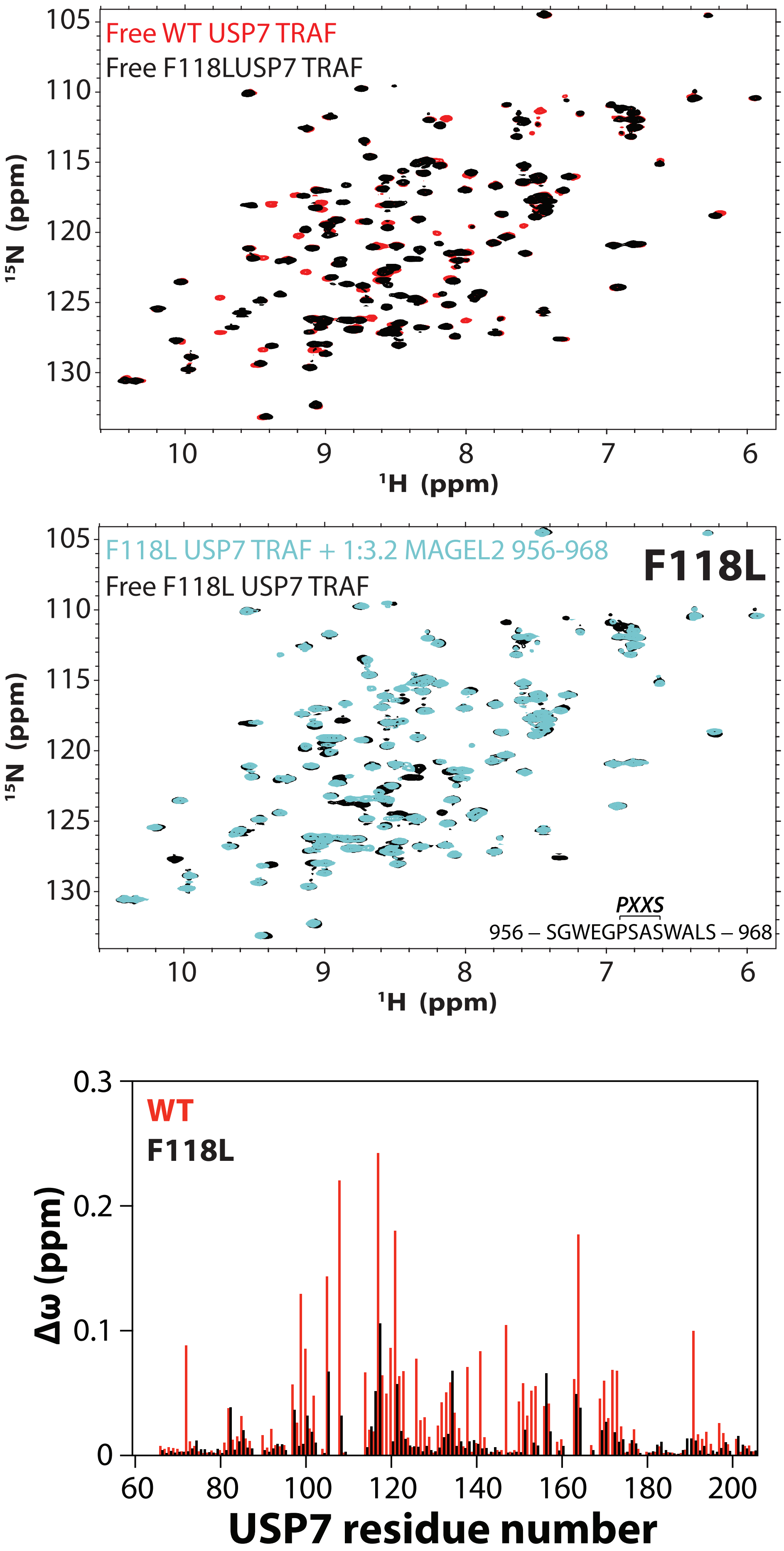
Free WT USP7 TRAF
Free F118LUSP7 TRAF
15N (ppm)
1H (ppm)
105
110
115
120
125
130
10
9
8
7
6
F118L USP7 TRAF + 1:3.2 MAGEL2 956-968
Free F118L USP7 TRAF
F118L
PXXS
956 – SGWEGPSASWALS – 968
WT
F118L
Δω (ppm)
0.3
0.2
0.1
0
60
80
100
120
140
160
180
200
USP7 residue number

### Figure S2

## C

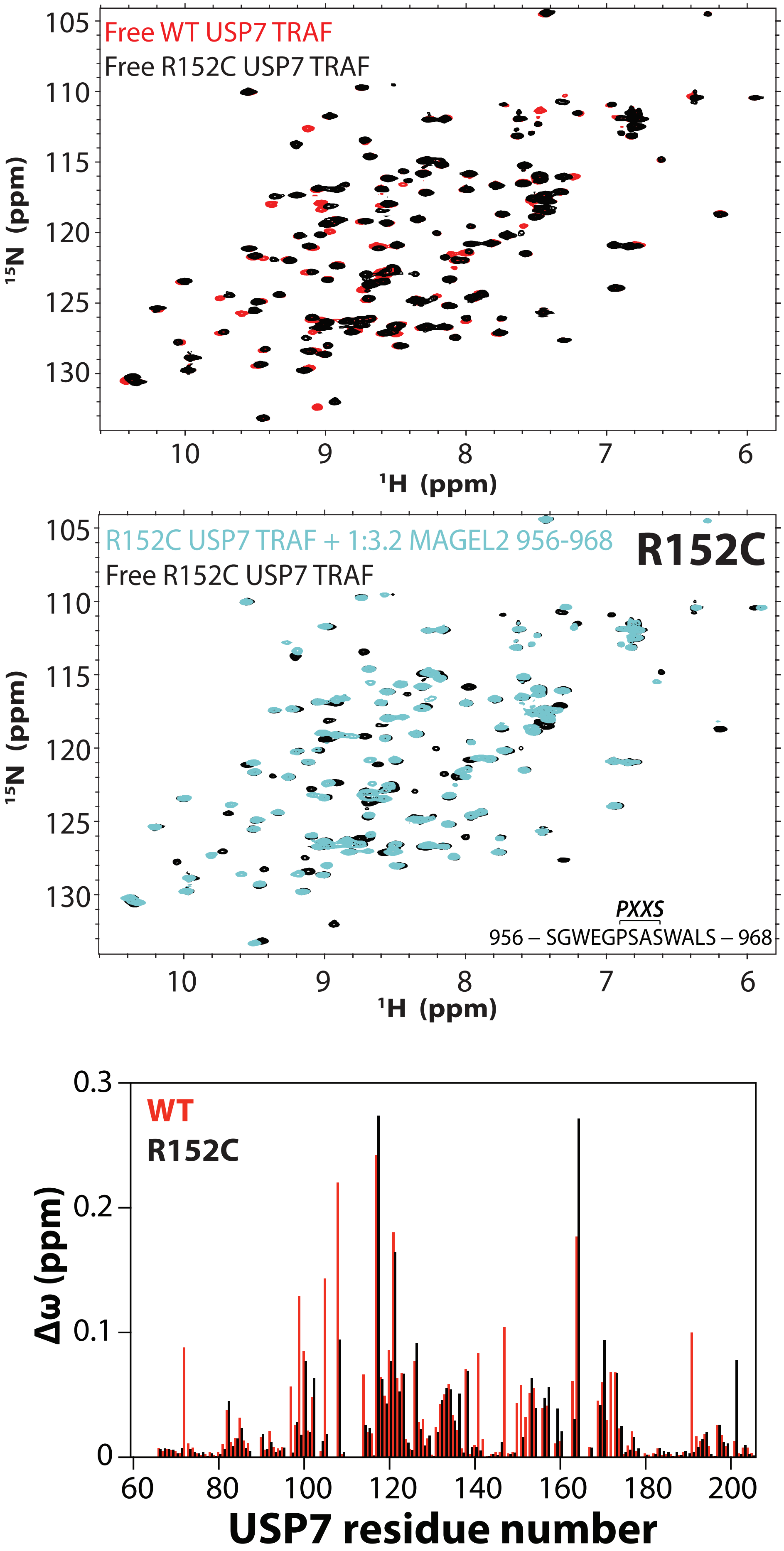

Free WT USP7 TRAF
Free R152C USP7 TRAF
105
110
115
120
125
130
15N (ppm)
10
9
8
7
6
1H (ppm)
R152C USP7 TRAF + 1:3.2 MAGEL2 956-968
Free R152C USP7 TRAF
R152C
PXXS
956 – SGWEGPSASWALS – 968
WT
R152C
0.3
0.2
0.1
0
Δω (ppm)
60
80
100
120
140
160
180
200
USP7 residue number

**Figure S2. NMR analysis of the effect of Hao-Fountain syndrome-linked TRAF domain mutations on MAGEL2 binding.**

Overlays of $^{15}N$-$^{1}H$ HSQC spectra of the WT TRAF domain and its variants H111Q (***A***), F118L (***B***), and R152C (***C***) are shown in the **top panels** and reveal minimal spectral perturbations, indicating that these mutations do not significantly alter the TRAF domain structure. The **middle panels** show overlays of the free (black) and MAGEL2-bound (blue) variants recorded in the presence of a 3.2-fold molar excess of unlabeled MAGEL2 peptide 956-968, revealing chemical shift perturbations consistent with binding. Per-residue NMR chemical shift perturbations ($\Delta\omega$) are quantified and shown in the **bottom panels** as bar plots (black bars). Comparison with $\Delta\omega$ values observed for WT TRAF binding to the same MAGEL2 peptide under identical conditions (red) reveals a significantly reduced binding affinity for the F118L variant.

Figure S3

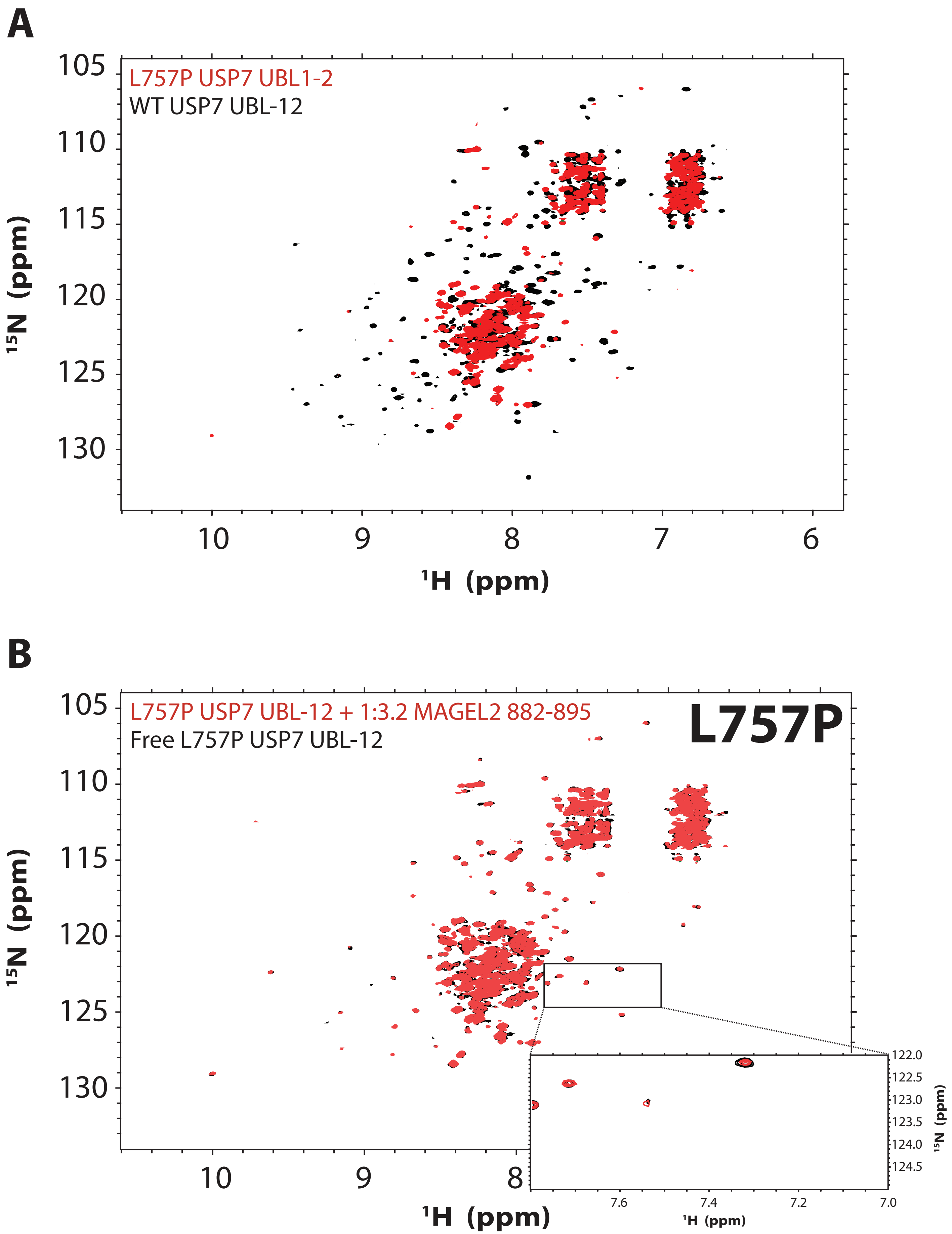

A
L757P USP7 UBL1-2
WT USP7 UBL-12
105
110
115
120
125
130
15N (ppm)
10
9
8
7
6
1H (ppm)
B
L757P USP7 UBL-12 + 1:3.2 MAGEL2 882-895
Free L757P USP7 UBL-12
L757P
105
110
115
120
125
130
15N (ppm)
10
9
8
1H (ppm)
122.0
122.5
123.0
123.5
124.0
124.5
15N (ppm)
7.6
7.4
7.2
7.0
1H (ppm)

**Figure S3. Hao-Fountain syndrome-linked variant L757P destabilizes the USP7 UBL1-2 domains.**

**A.** Overlay of $^{15}N$-$^{1}H$ TROSY spectra of the WT (black) and L757P UBL1-2 domains (red). The L757P mutation markedly reduces spectral quality, resulting in severe line broadening, loss of signal intensity, and an apparent reduction in spectral dispersion due to preferential loss of well-dispersed resonances. This behavior is consistent with aggregation and structural destabilization of the L757P variant compared to the WT. **B**. Addition of MAGEL2 peptide 882-895 at a 3.2-fold molar excess to the L757P variant does not restore spectral quality or induce detectable spectral changes (inset), consistent with severely impaired or abolished binding to MAGEL2.
